# High-powered, longitudinal mapping reveals genotype-phenotype stability and an unpredictable genetic architecture of adaptation

**DOI:** 10.1101/2025.11.04.686622

**Authors:** J.A. Smiley-Rhodes, M.C. Bitter, S. Berardi, J. Beltz, D.A. Petrov, P. Schmidt

## Abstract

The extent to which adaptation can be predicted is unknown. Here, we leveraged a longitudinal sampling design to test the efficacy of genomic prediction of trait evolution in an ecologically-relevant setting. Specifically, we monitored genome-wide allele frequencies and pigmentation variation in genetically diverse populations of *Drosophila melanogaster* across seven generations of evolution in both field mesocosms and a controlled, lab-based setting. At two points during trait evolution, we conducted a high-powered quantification of trait architecture that produced a well-resolved genotype-phenotype map. While we were able to use this map to correctly infer the direction of pigmentation evolution in both the field and lab mesocosms, the particular loci responding to selection, and thus the architecture of adaptation itself, was largely unpredictable. Further, we quantified a striking stability of the genotype-phenotype map, even across independent and genetically diverged populations. Our results hold implications for both the promise and limitations of genomic prediction.

## Introduction

Whether or not adaptation can be predicted is an outstanding question in evolutionary biology ^1,2^. This fundamental question permeates numerous issues of societal concern: from the preservation of species in the face of global change to the management of disease outbreaks. Two distinct ways in which adaptation can be predicted are: i) correctly inferring whether or not a focal trait was under selection, given patterns of genomic variation alone; or ii) correctly inferring the presence and direction of selection on genomic loci, given observations of phenotypic change alone. In order to employ either method of prediction, researchers must first quantify the molecular genetic basis of phenotypic variation and then understand how such phenotypic variation is translated into fitness-relevant differences among genotypes.

Instances in which adaptation has proven to be predictable are almost exclusively in the context of traits exhibiting simple genetic bases, whereby trait variation is explained by one or a few large effect loci ^3–9^. While such case studies provide fundamental insight into the evolutionary process, the vast majority of fitness-relevant traits instead exhibit complex genetic bases, whereby phenotypic differences are due to genetic variation at tens to thousands of loci spread throughout the genome ^10,11^. Accordingly, numerous recent studies have conducted genome-wide association studies of focal complex traits and then leveraged the resulting genotype-phenotype data to infer or predict trait evolution in natural populations^12–16^. Such efforts, however, may be confounded by the nature of complex trait variation itself.

Three facets of complex trait variation that may challenge evolutionary predictions are pleiotropy, epistasis, and genotype-by-environment (GxE) interactions. Pleiotropy occurs when a single genomic locus impacts phenotypic variation in multiple, fitness-relevant traits ^17,18^. As populations adapt, correlated selection across these traits can then impact the evolution of pleiotropic loci, constraining their effect on the evolution of the focal trait in question ^19^. In such a scenario, genetic variation at a mapped locus may behave as effectively neutral during the evolution of the trait itself, or even indicate trait shifts in the opposite direction of how it actually evolved through time ^10,18,20,21^. While pleiotropy complicates predictions through cross-trait correlation, epistasis – the interaction of allelic effects at multiple loci on a single trait – may shift the genotype-phenotype map itself throughout the evolutionary process ^22–25^. Specifically, epistasis can cause the marginal effect of trait-associated loci to change as a function of genetic background, which varies across populations with distinct demographic histories and patterns of selection ^22,23,25–33^. Compounding the effects of these forces are genotype by environment (GxE) interactions, which occur when the phenotypic effect of a genetic variant changes depending upon environmental exposure^30,34–36^. GxE can then cause the effect size of a locus mapped within a specific environmental context to be different from the effect of that locus on trait variation under the conditions in which selection acts. In concert, these facets of complex trait variation make it unclear whether loci mapped within a single population, environment, and point in time will be those that drive trait evolution in natural populations.

One novel way to systematically quantify the effects of pleiotropy, epistasis, and GxE on evolutionary prediction is through a longitudinal sampling design. Specifically, such an approach would involve constructing dynamical mapping data for a focal trait as it evolves in a natural setting, while simultaneously quantifying selection on genome-wide variation ^37–41^. With such data, the predictability of adaptation may be probed in a systematic manner, answering fundamental questions such as: are the loci underpinning phenotypic variation in a complex trait the same that respond to selection and underpin trait evolution in nature? How stable is the trait genotype-to-phenotype map throughout evolution and across different environments of selection? We took such an approach here, conducting a longitudinal analysis of trait variation and evolution of abdominal pigmentation in *Drosophila melanogaster*. We selected pigmentation as our focal complex trait because of its strong association with phenotypic diversity across the tree of life and implication in adaptive evolution across a range of taxa ^42^. For *D. melanogaster* in particular, abdominal pigmentation exhibits latitudinal clines among populations and rapidly and cyclically evolves in response to seasonally varying selection pressures ^43^. Importantly, abdominal pigmentation has long served as a model complex trait, whereby the biosynthetic pathway of the two main molecules, darkly pigmented melanin and light-colored sclerotin, have been well-characterized ^44,45^. More recent genome-wide association studies have pinpointed three core genes in the melanin/sclerotin biosynthetic pathway that explain the majority of intra- and inter-population pigmentation variation: *bric-a-brac* (*bab*), *tan*, and *ebony* ^46–48^. Interestingly, clines in pigmentation across space and time do not co-occur with expected clinal signatures of selection at the core pigmentation-associated loci ^43^, suggesting forces such as those discussed above may indeed affect trait variation and evolution across the species’ range.

Here, using large, genetically diverse replicate populations, we mapped pigmentation throughout evolution in mesocosms housed in two environments with distinct selective pressures: a simple, lab-based environment and a dynamic, semi-natural outdoor setting.^49^ The genotype-phenotype maps we constructed throughout evolution in each environment both validated the role of previously identified candidate pigmentation genes and also identified novel loci underpinning pigmentation variation in the species. We found that this map could then be used to correctly infer pigmentation evolution through time in each environment, given patterns of allele frequency data alone. Interestingly, however, the loci displaying signatures of selection as pigmentation itself evolved (i.e., the genetic architecture of adaptation) did not always behave in a manner as predicted by mapping, with some loci even shifting in the opposite direction from what might be predicted from the phenotypic shifts in the focal trait. This dynamic was exacerbated in the more complex, outdoor environment. Finally, we observed genotype-phenotype mapping to be remarkably stable and transferable across independent mapping populations, indicating that the marginal effects of mapped loci remain relatively stable even amidst shifting epistatic landscapes associated with the evolutionary process itself.

## Results

### Study scheme and power

Our study population was derived via outcrossing a panel of 76 inbred lines originally collected in local Pennsylvania orchards^50^. Several key advantages arise from this methodological decision. By allowing evolution to proceed from a single founding population we could evaluate the effect of epistasis on genotype-phenotype map stability directly and in a manner that is not confounded by population-specific demographic histories and patterns of linkage ^28,30^. Furthermore, the pigmentation and fitness-relevant variation subject to selection in our study was evaluated in the same geographic region where it is maintained naturally and in which we have documented repeatable seasonal evolution of lighter pigmentation from late spring through summer ^43^. Finally, the haplotype-informed allele frequency estimates we generated from our reconstituted outbred populations are of the accuracy produced using standard pooled sequencing approaches and depths ∼200x ^51^. Importantly, haplotype frequencies of samples collected throughout the experiment indicated that our reconstituted outbred population was not dominated by a subset of founding haplotypes, nor did we observe extinction of any founding lines throughout the progression of evolution in our mesocosms (Fig. S1-2).

The reconstituted outbred population was used to seed each of 10 indoor cage replicates maintained in controlled, lab-based conditions, as well as 10 replicate outdoor mesocosms exposed to natural environmental fluctuations in Philadelphia, PA (Fig. 1) ^49,52,53^. This paired design was leveraged to gain mechanistic insights into the primary drivers of pigmentation evolution observed in natural populations – specifically, whether the trait is under selection elicited by fluctuating abiotic variables such as temperature, or also shaped by the biotic stressors (namely rapidly increasing density) shared between our study environments. Selection in this latter case could be direct, or indirect, due to shared genetic correlations of pigmentation with the reproductive traits that come under selection during increasing population densities^49,54^. Over the course of ten weeks and approximately seven generations of evolution, we quantified phenotypic evidence of direct (or indirect) selection on pigmentation and monitored genome-wide frequency changes of 1.9 M single nucleotide polymorphisms (SNPs). At two time points throughout this process (weeks 4 and 10), and in each environment and replicate cage, we conducted a tail-based mapping: pools of 75 female flies, each representing the 15% lightest, 15% darkest, and representative midpoint of the trait value distribution (hereafter referred to as ‘light’, ‘dark’, and ‘midpoint’ trait value groups, see Methods) were isolated and sequenced to identify pigmentation- associated SNPs. We used simulations to evaluate the statistical power of our tail-based mapping approach, specifically its ability to identify quantitative trait loci over a range of effect sizes and starting frequencies (see Methods). Congruent with recent work^55^, we found that our highly-replicated, tail-based mapping exhibits power to detect complex trait loci comparable to an inbred line genome-wide association study using on the order of 800 – 900 strains (Fig. S3-4)^48^.

**Figure 1.**
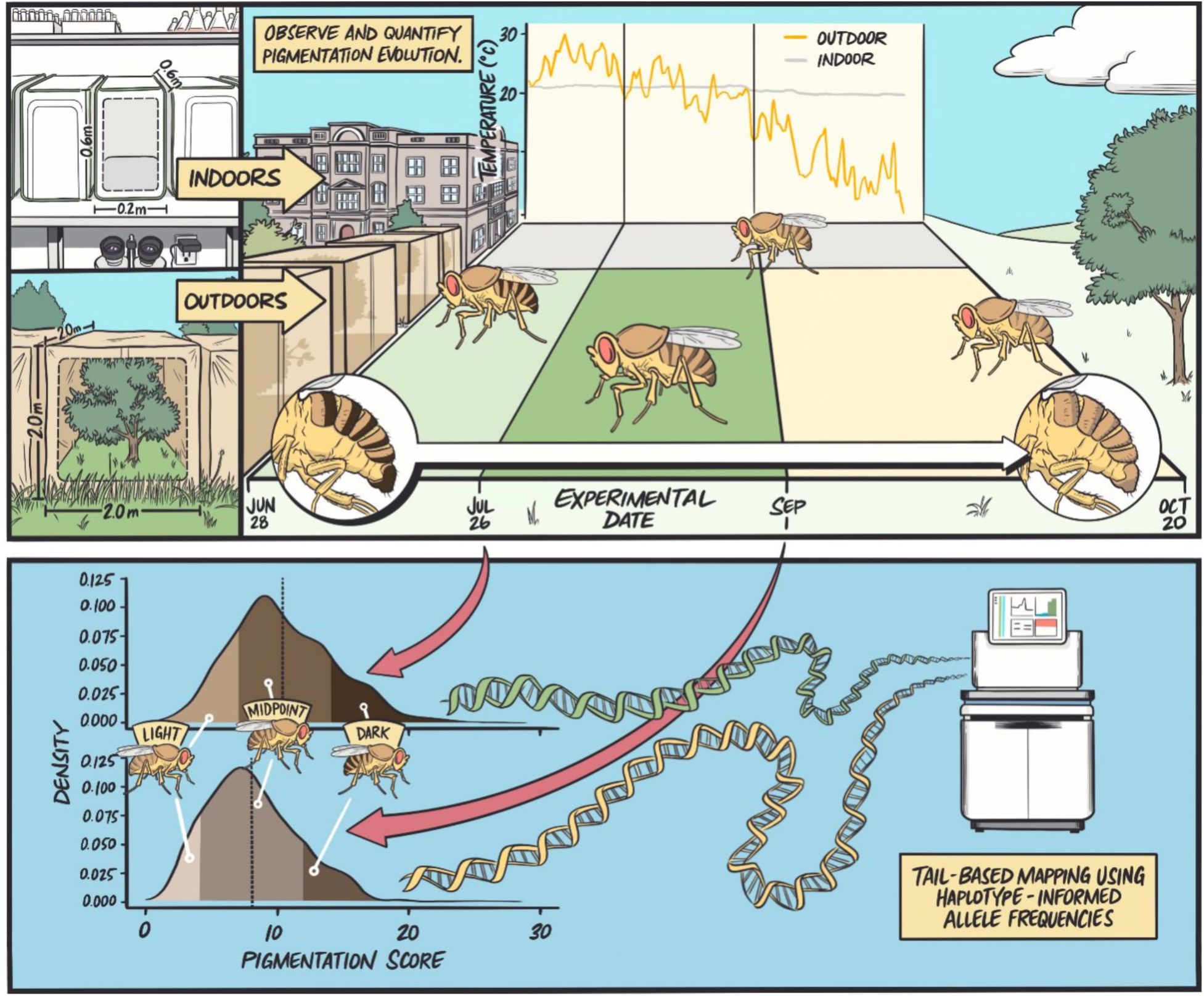
Experimental design. Ten cages each in an indoor laboratory and in outdoor, field mesocosms were seeded with an outbred population of *D. melanogaster*. The replicate populations were allowed to evolve from late June to late October, during which time significant pigmentation evolution was observed in each environment (see text). Four and ten weeks after the seeding of cage replicates, we conducted tail-based mapping of pigmentation. Specifically, following two generations of common garden rearing, flies were segregated into color fractions based on the population’s distribution of abdominal pigmentation scores (fractions comprised 75 female individuals from the darkest 15%, the lightest 15%, and a representative sample from the trait midpoint). Each color fraction from each replicate cage, environment, and time point was then used in whole-genome sequencing and estimation of high-accuracy, haplotype-informed allele frequencies at 1.9 M SNPs.

### Phenotypic and genomic variation across environments and throughout evolution

We first determined whether pigmentation evolved in our experimental populations and whether this was driven by natural selection. Specifically, we quantified pigmentation independently for each inbred reference line, across individuals in the derived outbred population (hereafter, ‘baseline’), and among replicate populations at weeks four and ten of evolution in each environment. As expected, our baseline population exhibited variation in pigmentation that was representative of the variation observed across the 76 inbred lines from which it was derived (Fig. 2A). Pigmentation after two generations of evolution resulted in significantly lighter pigmentation across replicates in both the lab and field environments (pairwise t-tests, week 4 outdoor p-value << 0.001, week 4 indoor p-value << 0.001), and the replicate populations then evolved in a parallel manner to become significantly lighter during the six additional weeks of evolution in both the outdoor and indoor environments (pairwise t-tests; week 4 vs. 10 outdoor p-value << 0.001, week 4 vs. 10 indoor = 0.0245). Notably, a significant treatment by time interaction on variation in pigmentation throughout the experiment was observed (two-way ANOVA; F = 17.03, p-value << 0.001), driven by the accelerated rate and magnitude of lighter pigmentation evolution in the outdoor mesocosms (Fig. 2A).

**Figure 2.**
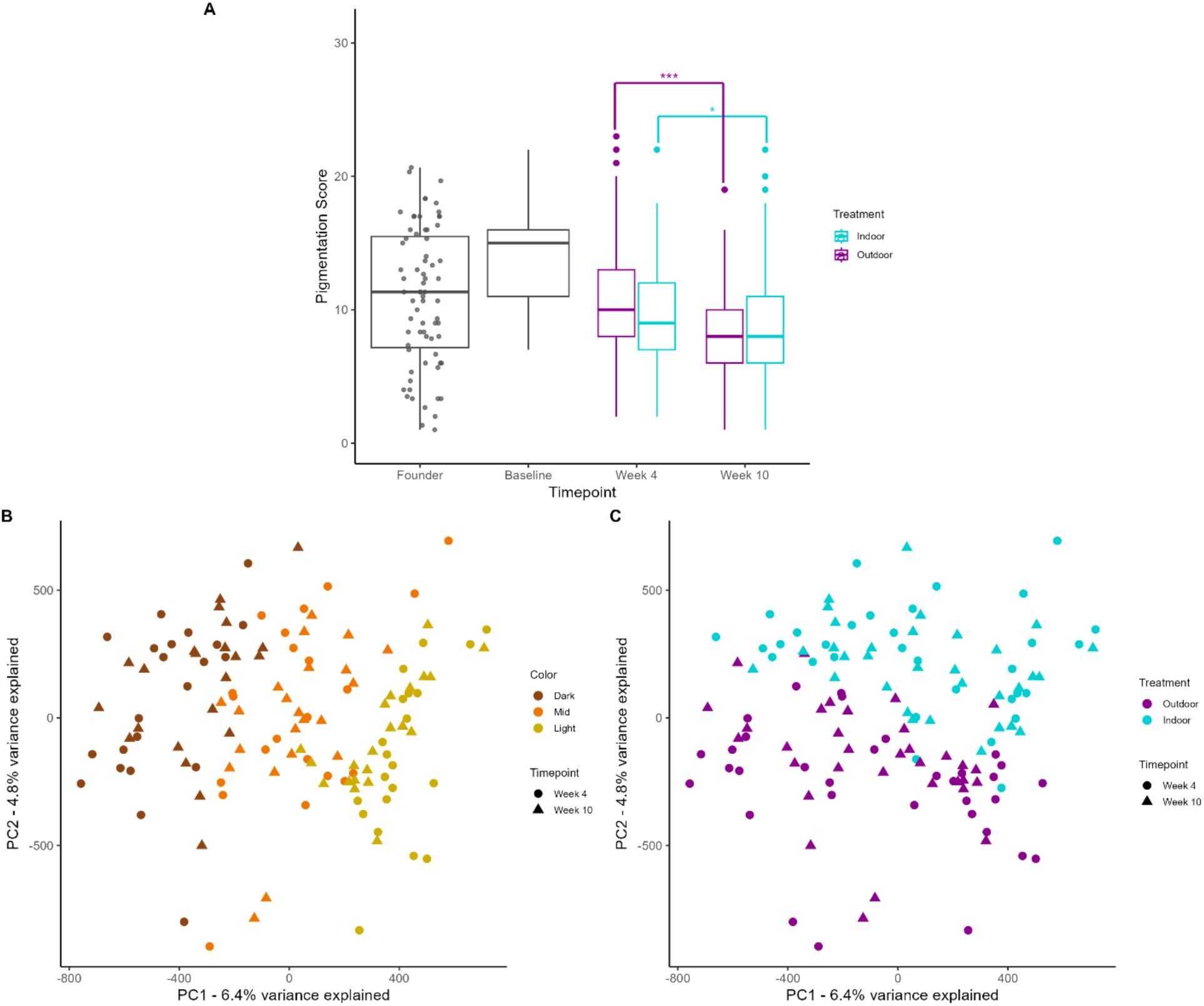
Phenotypic and genomic variation throughout study period. (A) Distribution of the total pigmentation scores (whereby higher scores indicate darker pigmentation) across samples. “Founder” refers to our 76 inbred lines that we outbred for four generations to generate our baseline (week 0) population. Grey dots each represent the average pigmentation score of N = 3 females from each inbred line. The baseline pigmentation distribution represents the scores of N = 90 females from the initial outbred population which was used to seed our indoor and outdoor cages. From the samples collected during the experiment at weeks 4 and 10 in both environments, we scored 30 females per replicate cage (N = 300 in total for each timepoint/treatment combination) and found a significant shift towards lighter pigmentation in both outdoor and indoor populations (pairwise t-test; p-values << 0.001). (B-C) Projection of samples used in trait mapping onto the first two principal components generated via PCA of allele frequency data from all 1.9 M SNPs. In (B) samples are colored by the color fraction from which they originate (dark, midpoint, or light) and the shape of the points corresponds to the timepoint when the sample was collected. PC1 value and color fraction identity are significantly associated (linear regression p-value << 0.001). In (C) samples are colored by treatment. PC2 value and treatment are significantly associated (ANOVA p-value << 0.001).

Next, we determined whether there existed evidence of environment-specific patterns of selection on genomic variation that could, in effect, induce distinct epistatic landscapes between the lab and field-based mapping populations. Specifically, we ran principal component analysis (PCA; *prcomp* function, R) on the normalized allele frequency data of all 1.9 M SNPs and then projected samples onto the first two principal components (Fig. 2B-C). Strikingly, the first principal component (6.4% variance explained) segregated samples in accordance with trait group (Fig. S5, R^2^ = 0.762; p-value << 0.001), while the second principal component (4.8% variance explained) segregated samples by environment of selection (ANOVA p-value << 0.001). These patterns largely held when PCA was conducted separately for each chromosomal arm (Fig. S6-7). This analysis thus demonstrates the pronounced impact of both phenotypic group and environment of selection on genome wide patterns of variation, while also suggesting that a shared set of loci underpin variation in pigmentation across mapping populations.

### Ancestry-informed mapping provides resolved architecture of abdominal pigmentation

We next aimed to map the underlying genetic architecture of pigmentation. We initially focused mapping on samples collected at week 10 in the outdoor mesocosms, as these replicate populations had undergone evolution in a semi-natural environment and experienced more generations of recombination since initial outbreeding than the week 4 samples (and were thus predicted to generate better-resolved genetic maps). Our general approach to mapping was to identify SNPs harboring alleles with systematic frequency changes across the light, midpoint, and dark trait value groups. However, given the finite number of haplotypes from the founding inbred lines used to derive our outbred population, a significant association between a SNP and trait value group could indicate either its linkage to a locally causal allele and/or its broader inbred-line ancestry. Accordingly, to determine the magnitude of this ancestry effect genome-wide, we quantified the average contribution of haplotypes coming from each inbred line to each trait value group and tested for the association of this quantity with the pigmentation score of that inbred line. The results of this are depicted in Figure 3A, where we observe that the light trait value groups across replicates contain a disproportionate amount of ancestry from the lightest inbred lines and, likewise, the dark trait value groups contain a similarly disproportionate amount of ancestry from the darkest inbred lines (R^2^ = 0.032; p < 0.01). To further quantify this ancestry effect on pigmentation, we ran a simple regression (GLM; logistic link function and quasibinomial error variance) of allele frequencies on trait value group. We took the top 1% of the SNPs (based on nominal p-value from the GLM) and calculated their expected frequency difference between light and dark color fractions based on estimates of average contribution of each inbred line to that sample, as well as the association between that SNP and its presence in the inbred line (Fig 3B). Indeed, we see that those GLM-identified SNPs with a higher relative frequency in the light color fraction tend to be associated with the lighter pigmented lines, making it hard to disentangle the SNP’s local association with a causal locus from its association with global line ancestry.

**Figure 3.**
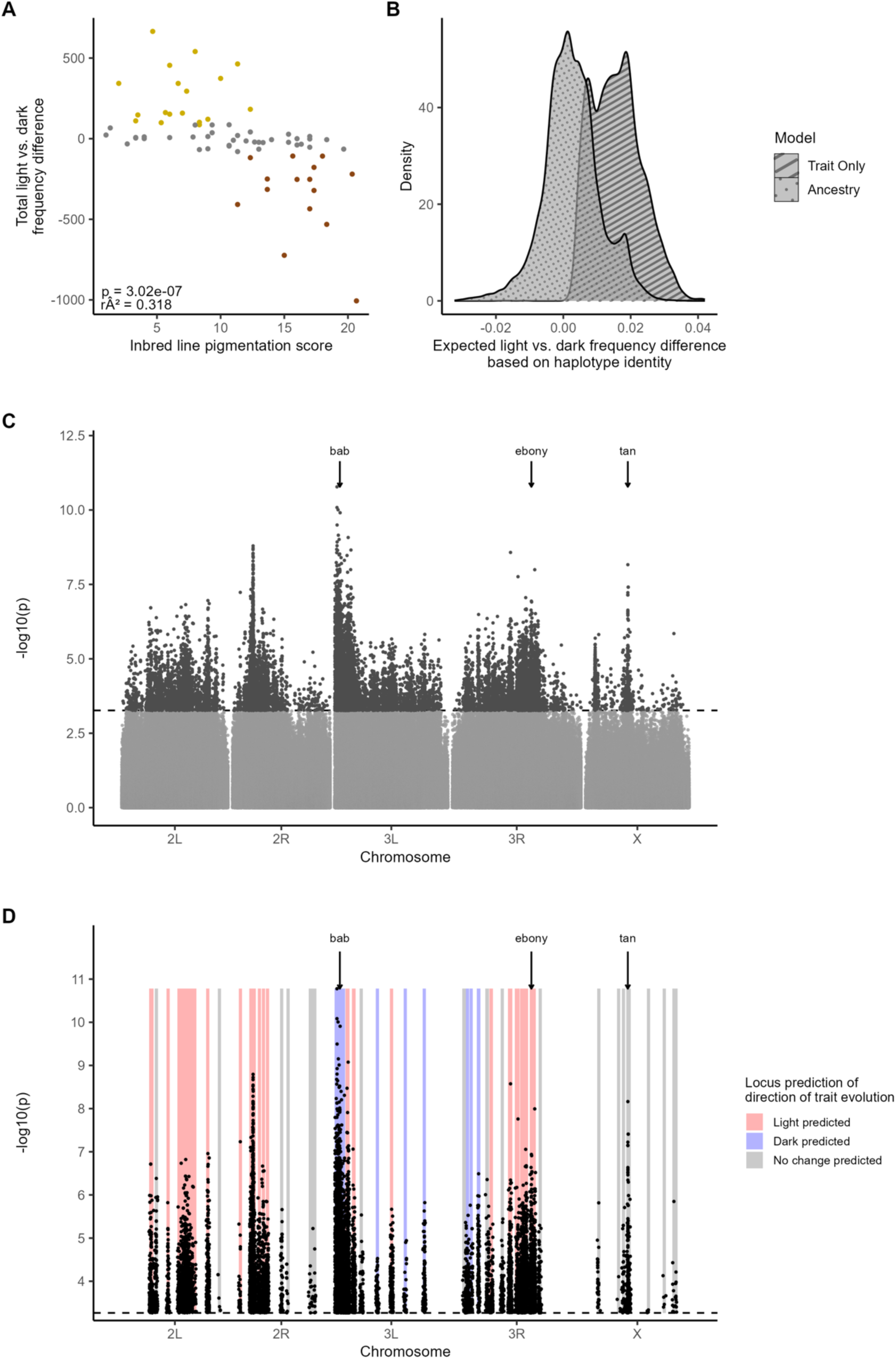
Genetic architecture of female abdominal pigmentation. (A) The pigmentation score of the founding inbred lines compared with the light vs. dark frequency differences across all samples for the haplotypes originating from these lines. Inbred line pigmentation is highly correlated to the haplotype’s light vs. dark enrichment (linear regression p-value << 0.001). Colored points indicate inbred lines that drive the significance of the relationship between pigmentation score and enrichment in color fractions (gold points correspond to inbred lines enriched in light color fraction and dark brown to lines enriched in dark color fractions) (B) Distribution of the difference between average expected light frequencies and average expected dark frequencies for the top 1% of SNPs from a model only considering the impact of trait value on allele frequency variation (Trait-Only), versus a model with an additional term that represents the expected frequency in the sample based on the SNPs’ ancestry (Ancestry). (C) Manhattan plot depicting nominal p-values derived from an ancestry-informed mapping of SNP association with pigmentation variation. The dashed horizontal line corresponds to an empirical FDR threshold of 0.0005, derived via permutations. (D) Genomic distribution of SNPs confined within N = 42 unlinked, independent loci underpinning pigmentation variation. SNPs are plotted as a function of their genomic coordinate and regression-derived nominal p-value (as in (C)). Shaded regions denote the start and end points of each of the identified unlinked, independent loci. Shaded region color then denotes the predominant behavior of light-associated SNPs within each locus through time: red shading indicates loci with light-associated alleles favored throughout evolution of lighter pigmentation (18 total loci; binomial test, FDR < 0.05), blue shading indicates loci in which the light-associated alleles are predominately selected against (7 total loci; binomial test, FDR < 0.05), and grey shading indicates loci with no signature of selection on light-associated alleles (17 total loci; binomial test, FDR > 0.05). The location of the canonical pigmentation genes, *bric-a-brac (bab), ebony,* and *tan*, are indicated on the Manhattan plots. Potential candidate genes underlying the additional loci observed throughout the genome are discussed in the Supplement.

To account for the strong ancestry effects on SNP-pigmentation association, we modelled allele frequency as a function of both pigmentation trait value and an ancestry-predicted frequency term – the latter capturing each SNP’s expected frequency given its association with each inbred line and that line’s genome-wide relative frequency across the sample (model formula: allele frequency ∼ trait value + ancestry-predicted frequency). Thus, we controlled for SNPs that had an elevated light-dark frequency difference that could be entirely explained by linkage to high-frequency haplotypes. As expected, in Fig. 3B we see that the distribution of expected light-dark frequency differences for SNPs identified by this model is now centered around zero, indicating that the global bias produced by our trait-only GLM is eliminated. We further validated that SNPs identified via this approach were not exclusive to the lightest nor darkest founding inbred lines (colored points, Fig. 3A), but rather present at appreciable frequency across all founding haplotypes (Fig. S8). We defined a robust set of pigmentation-associated SNPs by permuting trait values and re-running our ancestry-informed GLM (N = 100 permutations). A SNP was considered significant if its observed p-value exceeded the 99.95^th^ percentile of the permuted p-value distribution (corresponding to an empirical FDR threshold < 0.0005; Fig. S9). Even at this conservative threshold, we see that trait-associated SNPs are pervasive throughout the genome and distributed across all chromosomal arms (Fig 3C). Hereafter, SNPs identified via this approach are referred to as either light- or dark-associated in accordance with their relative frequency in the light and dark color fractions (e.g., the light-associated allele for each mapped SNP is that occurring at higher relative frequency in the light color fraction).

We next used our set of pigmentation-associated SNPs to identify the number and location of unlinked loci underpinning variation in the trait. Our procedure to do so was as follows: we first identified the most significant SNP on each chromosomal arm, as well as its surrounding, locally un-recombined region (defined by the recombination rate and number of generations since cage founding). We then eliminated all trait-associated SNPs on the chromosome whose relative frequency in the light and dark fractions could be explained by haplotype frequencies within the locally un-recombined focal region alone. In effect, this eliminated all GLM signal due to linkage to the focal SNP. From the set of remaining SNPs, we identified the SNP with the most significant nominal p-value and repeated this procedure. This was iterated until we exhausted all trait-associated SNPs on the chromosome and resulted in a total of 76 genomic loci, the midpoint of which was the most significant, locus-defining SNP (see Methods and Fig. S10). Merging loci with overlapping start and endpoints, we obtained a final list of 42 loci, ranging in size from 700 Kb to 3 Mb, and distributed across all chromosomal arms (Supplementary Data File 1). This indicates that the genetic architecture of our focal, ecologically relevant complex trait, is not underpinned solely by a single or few large-effect loci, nor exclusively by a very large number of weak-effect loci (e.g., 1000’s). Rather, we infer the presence of a moderate number of large to moderate-effect loci underpinning trait variation, while acknowledging there may be further loci of small effect that our approach lacks sufficient power to detect. We validated that pigmentation-associated SNPs within the identified loci were not exclusive to the lightest and darkest haplotypes enriched in the trait tails (Fig. 3A), but rather shared across all founding inbred lines (Fig. S11).

We quantified the extent to which our identified loci validated previously described, and potentially identified new, candidate genes associated with pigmentation variation in the species. Reassuringly, the top loci on chromosomal arms 3L, 3R, and X contained the previously described core pigmentation genes: *bab*, *ebony*, and *tan* ^44,56–58^. A systematic comparison of the association signal between loci identified here and two recent, well-powered GWAS of the trait ^47,48^ indicated significantly more overlap than expected by chance (permutation p < 0.001, see Methods). However, this signal is eliminated when we no longer consider SNPs within the loci containing *bab*, *ebony*, or *tan* (p = 0.994), suggesting little overlap outside of these three core genes. We suspect the extensive enrichment of pigmentation-associated variation outside the canonical loci we observe to represent the discovery of novel pigmentation-associated loci, which likely arises both due to the exceptional power of our mapping (Fig. S3-4) and also potential differences in the underlying causal loci impacting pigmentation variation among the population used here and those in previous studies^59,60^. In the supplement, we discuss candidate genes within these novel loci that likely harbor pigmentation-relevant variation (Supplementary Discussion; Table S1-2) and below demonstrate, using out-of-sample phenotypic prediction, that these novel loci are unlikely to be false positives (see: *Stable genotype-phenotype mapping throughout evolution and across environments of selection*).

### Genomic predictions of complex trait evolution

One application of a well-resolved genotype-phenotype map is to predict whether or not a trait has evolved in a natural population using genomic data alone. Our longitudinal study design afforded the unique opportunity to test the efficacy of this approach directly by leveraging our well-resolved genetic mapping data (Fig. 3C-D) in concert with the time-series allele frequency data we collected throughout phenotypic evolution in the outdoor mesocosms (Fig. 1; 2A). Specifically, for each of the 42 unlinked loci identified above, we quantified the frequency shifts of the light-associated alleles over the period during which the outdoor replicate populations evolved to become significantly lighter (weeks 3 to 10; Fig. 2A). We then inferred whether the observed allele frequency movement within each locus indicated lighter pigmentation evolution by comparing the distribution of shifts at mapped alleles to a distribution derived from a set of matched control SNPs, which served as a proxy for the magnitude of allele frequency change expected as a result of genetic drift or sampling noise (binomial test; see Methods). We found that out of the 42 loci, 18 exhibited an overall increase in light-associated allele frequency and thus behavior through time concordant with what would be expected given the evolution of lighter pigmentation. However, for 17 mapped loci we did not detect any systematic shift indicative of trait evolution, while 7 loci showed frequency shifts indicative of the evolution of darker pigmentation (Fig. 3D). Thus, in sum, while our genome-wide inference correctly predicted evolution of lighter pigmentation in the outdoor mesocosms (p-value << 0.001; Fig. 4A) we observed that at particular loci and even entire chromosomal arms (Fig. S12), the genetic basis of pigmentation evolution was unpredictable based on mapping data alone. Strikingly, this unpredictable genetic architecture of adaptation was evident at our loci of largest effect: while allele frequency shifts within the *ebony* locus, as well as the largest effect locus on 2R, correctly predicted evolution of lighter pigmentation (light-associated alleles increased in frequency during the evolution of lighter pigmentation), variation within the locus containing *bab* predicted evolution of darker pigmentation (light-associated alleles decreased in frequency during lighter pigmentation evolution) and the *tan* locus behavior suggested that pigmentation did not evolve (light-associated alleles behaved neutrally during lighter pigmentation evolution) (Fig. 3D).

**Figure 4.**
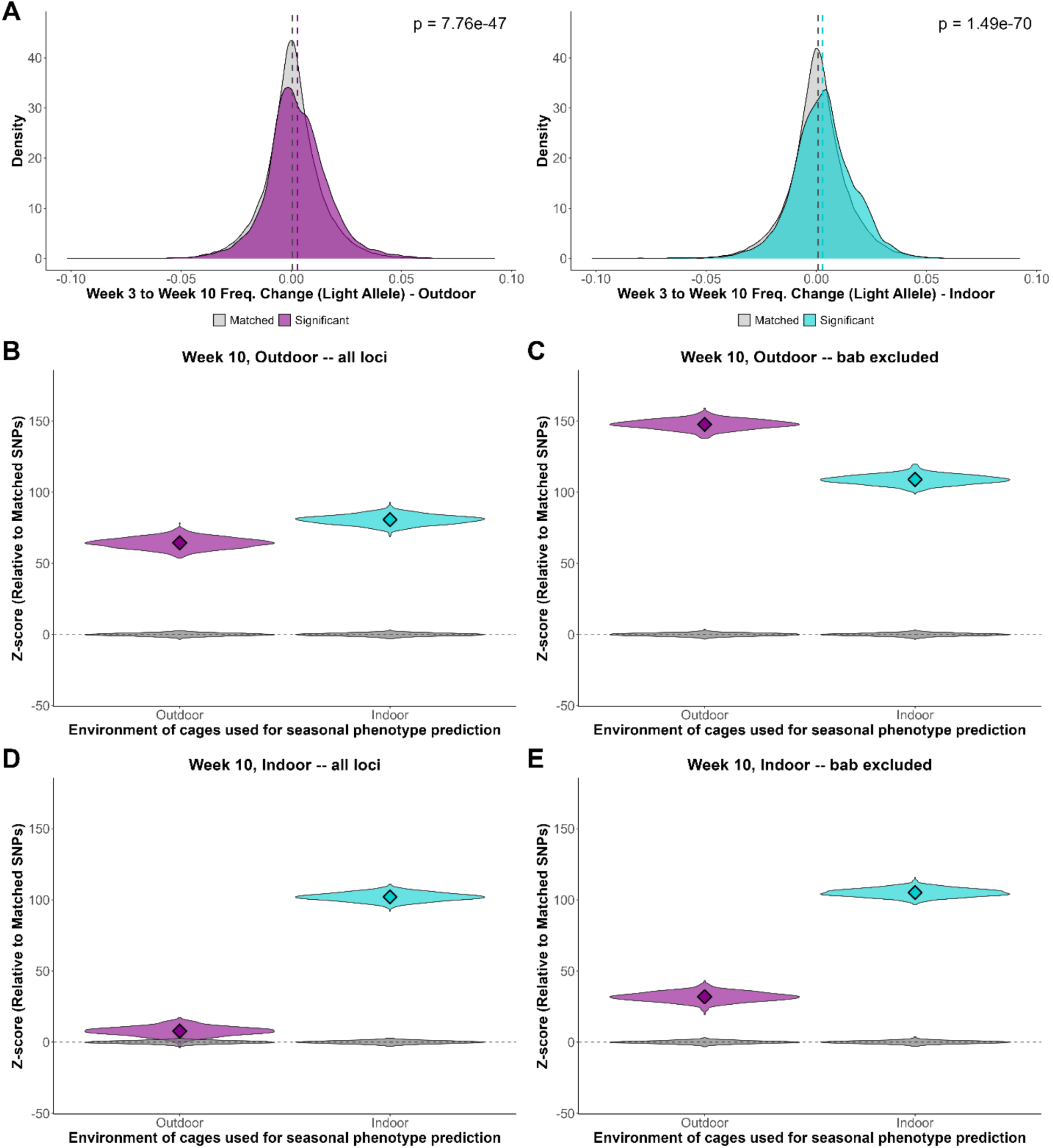
Prediction of phenotypic evolution in simple and complex environments. (A) Genome-wide allele frequency change at pigmentation-associated SNPs identified in the outdoor, week 10 mapping population and quantified throughout trait evolution in the outdoor (left, purple) and indoor (right, blue) mesocosms, relative to matched control shifts (grey distributions). (B) Distribution of Z-scores for bootstrapped estimates of genomic predictions for pigmentation evolution in the outdoor (magenta) and indoor (turquoise) mesocosms using SNPs identified in the outdoor, week 10 mesocosms. Positive Z-scores indicate net lighter pigmentation evolution while negative values would indicate net darker evolution. Z-scores were computed by comparing inferred relative phenotypic change using mapped SNPs to that computed using a set of matched control SNPs (grey distributions). (C) Same as in panel (B), but genomic predictions discard pigmentation-associated SNPs within the *bab* locus. (D-E) Same as (B-C) but for pigmentation-associated SNPs identified via mapping the week 10 indoor populations.

### Efficacy of genomic prediction of trait evolution differs between complex and simple environments

The unpredictable behavior of mapped loci during trait evolution we quantified above manifested in a field setting where replicate populations were exposed to a dynamic suite of fluctuating selection pressures. This raises the question of whether this unpredictability would be subdued in a simpler, lab-based environment with fewer selective pressures overall. Our study design provided a unique opportunity to test this hypothesis via comparing genomic predictions of trait evolution for the replicate populations evolved in the outdoor mesocosms relative to those that evolved within the indoor, lab-based mesocosms (Figs. 1-2). Accordingly, we first quantified whether light-associated alleles were favored during the evolution of lighter pigmentation in each environment, finding that their genome-wide frequencies increased significantly relative to background allele frequency change throughout trait evolution in both the lab and outdoor mesocosms (pairwise t-test p-values << 0.001; Fig. 4A; also see Fig. S12-S14). Next, we used the relative effect sizes produced by our mapping data, as well as their magnitude of allele frequency during trait evolution, to provide a genome-wide summary statistic of pigmentation evolution in each environment (Fig. 2A, Methods). This approach is analogous to polygenic score-based predictions of trait evolution oftentimes implemented in studies of human adaptation ^13,14,16^. Still, it is important to note that the effect size estimates of the mapped loci obtained here are derived from regression of group trait values on allele frequency data, rather than individual-level trait and genotype data; our phenotypic predictions then more generally represent a genomic prediction of relative (not absolute) trait value through time. Encouragingly, this approach correctly inferred that lighter pigmentation evolved in both the indoor and outdoor environment (Fig. 4B). Interestingly, the predicted magnitude of phenotypic change was greater in the indoor environment, despite our phenotypic assays showing that replicate populations in this environment evolved to a lesser degree (Fig 2A).

We next systematically evaluated the impact of the *bab* locus on the accuracy of our genomic prediction of trait evolution, as it is a canonical pigmentation gene, has documented pleiotropic effects across a range of fitness-relevant phenotypes^61–64^, and, in our experimental populations, harbors light-associated alleles that became selected against during lighter pigmentation evolution in the outdoor mesocosms (Fig. 3D). By excluding mapped alleles at the *bab* locus and reconducting our genomic prediction, we observed that the predicted magnitude of lighter pigmentation evolution increased in both environments and, strikingly, to a much greater degree in the outdoor environment (Fig 4C). This analysis thus provides a key example of how underlying pleiotropy (or GxE) at a single locus may skew genomic predictions of phenotypic evolution through time. In the supplement, we further explore how genomic predictions of trait evolution using only variation within the canonical pigmentation loci alone can lead to spurious predictions of trait evolution (Fig. S15-16).

The genomic predictions of trait evolution presented thus far were based on mapping data procured from the week 10, outdoor populations. Accordingly, we next quantified if and how these predictions differed when derived from mapping results generated using samples from week 10 in the indoor environment. Specifically, using the same mapping and prediction procedure, we found that while this independent set of mapped loci correctly predicts trait evolution in the indoor environment, it fails to predict evolution of lighter pigmentation in the outdoor environment (Fig. 4D). This discrepancy in genomic prediction of trait evolution in the outdoor environment (i.e., comparing magenta distributions in Fig. 4B vs. Fig. 4D) demonstrates that prediction in complex, natural environments may be hampered by using genetic maps generated in simple, lab-based environments, even though the populations are matched in every other way. In contrast, evolution in the simple, controlled indoor environment was correctly predicted, no matter the environment of the mapping population. Strikingly, but also consistent with those genomic predictions generated using the outdoor mapping data (Fig. 4B-C), we found that the exclusion of *bab* locus variation allowed us to correctly predict trait evolution in the outdoor environment using the indoor mapping data (Fig. 4E).

### Stable genotype-phenotype mapping throughout evolution and across environments of selection

How well do genotype-phenotype maps derived from a single population, at a single point in time, transfer to genetically diverged populations? This represents another fundamental question in complex trait genetics and evolution that we could address within the context of our longitudinal, multi-environment study. Specifically, we observed a pronounced signature of genome-wide divergence associated with the difference between our lab-based, indoor environment and the outdoor, field mesocosms (Fig. 2B), and we previously reported substantial evolutionary change within each of these environments throughout the observation period monitored in this study ^49,52^. One implication of this environmental and temporal genomic divergence is that it may induce shifts in the epistatic landscape and/or elicit context-dependent dominance of pigmentation-associated loci. In effect, these processes may alter the marginal effect of pigmentation-associated loci and generate distinct genotype-phenotype maps that preclude phenotypic prediction across mapping populations ^30,65,66^.

We evaluated the stability of the genotype-phenotype map by re-implementing our mapping approach, as described above, on the replicate populations sampled in outdoor mesocosms at week 4, as well as the indoor populations sampled at weeks 4 and 10. The number of loci underpinning variation across all mapping populations was comparable (ranging from 30-43 independent loci, Supplementary Data File 1; Table S2). Furthermore, as expected, the total genomic distance spanned by these loci declined from weeks 4 to 10 (from ∼1 Mb at week 4 to ∼700 Kb at week 10) due to additional recombination events that allowed for more localized mapping.

The similarities in architecture inferred in the lab and field environments at week 10 are clearly apparent in the mirrored Manhattan plots presented in Fig. 5A, where regions of heightened pigmentation-associated variation are replicable across the week 10 outdoor and indoor mapping populations. Notably, loci identified across all four mapping populations contained the canonical pigmentation genes *tan*, *ebony*, and *bab* (Fig. S17), demonstrating consistency of mapping as well as the pronounced effect these genes have on pigmentation variation in this species. We next took two approaches to rigorously quantify, genome-wide, the similarity in loci harboring pigmentation-associated variation across the four independent mapping populations. First, we assessed whether the start and end points of loci identified in each independent mapping population were closer than expected by chance: for each set of loci identified within a particular mapping population, we selected the locus-defining, midpoint SNP (i.e., that with the most significant p-value) and then quantified the number of times it fell within a locus in the other three sets of mapped loci (each locus could therefore overlap between 0 to 3 times with the remaining mapping populations). The number of overlaps across all loci for a particular mapping population were then summed and compared to an empirical null distribution derived by iterating this procedure N = 2000 times using a set of randomly selected SNPs (see Methods). Across sets of mapped loci, midpoint SNPs were far more likely to fall within a locus of an independent mapping than expected by chance (Fig. 5B), demonstrating that the regions harboring pigmentation-associated variation were conserved across mapping populations. To probe this consistency further, we quantified the distance between locus midpoints among mapping populations. As expected, these distances were significantly closer than expected based on an empirical null (Fig. 5C, see Methods), further indicating the conservation of pigmentation-associated loci across mapping populations.

**Figure 5.**
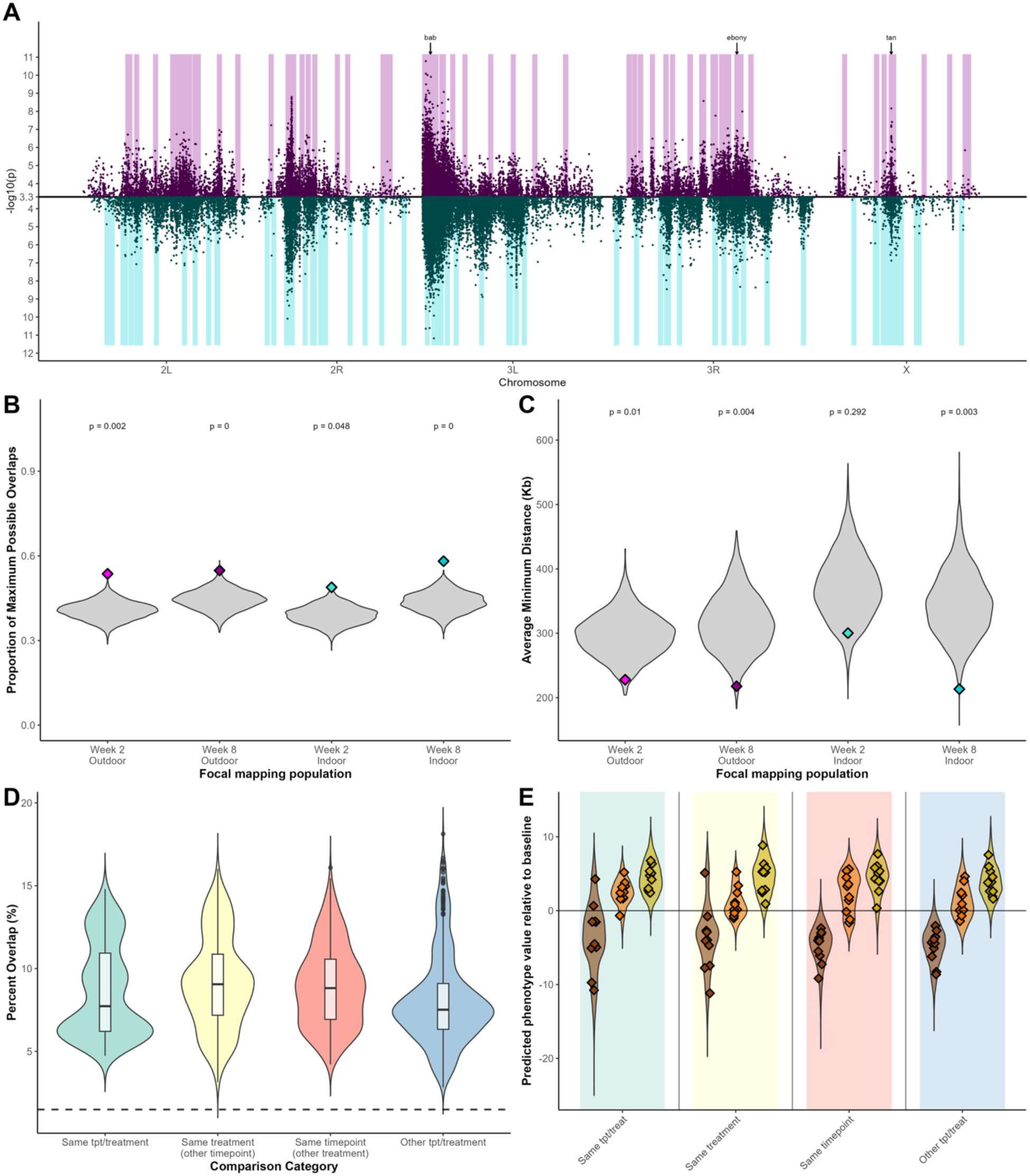
Epistasis does not preclude portability of trait mapping. (A) Mirrored Manhattan plots of genetic architecture quantified at week 10 in the outdoor (top) and indoor (bottom) environments. Only SNPs with a p-value corresponding to empirical FDR < 0.0005 are depicted. For each mapping, shaded regions correspond to unlinked loci underpinning pigmentation-associated variation. (B) Observed (diamonds) vs expected (grey violin distributions) overlap between pigmentation-associated loci across mapping populations. (C) Observed (diamonds) vs expected (grey violin distributions) distances between midpoint SNPs of trait associated loci between mapping populations. (D) Observed percent overlap of trait-associated SNPs identified within and between mapping populations. (E) Predicted relative phenotypic values for dark (brown points), midpoint (orange points), and light (yellow points) color fractions using genotype-phenotype mapping conducted in the outdoor mesocosms at week 10 where positive and negative values indicate lighter or darker pigmentation relative to the baseline population, respectively. Predictions (diamonds) were conducted for each trait value group and replicate cage. Colored violins represent 1000 bootstrapped predictions of each predicted, relative phenotypic value.

Given the pronounced mapping stability at the locus-level, we next interrogated whether pigmentation-associated SNPs were also shared across mapping populations. To do so, we reidentified trait-associated SNPs using our ancestry-informed GLM in groups of 5 (of the 10 total) replicates per mapping population. This then allowed for a balanced comparison of the overlap in trait-associated SNPs within subsets of the same mapping population (e.g., overlap in trait associated SNPs between independent subsets of five replicates from the outdoor week 10 samples), and between mapping populations (e.g., overlap in trait associated SNPs between subsets of five outdoor, week 10 and five indoor, week 4 replicates). Strikingly, we found that the amount of overlap in SNPs identified between subsets of the same mapping population did not exceed that between different mapping populations, either from the same environment and a different time point or even between environments (Fig. 5D; Fig. S18). In the supplement, we further quantified this SNP-level parallelism using a leave-one-out cross validation scheme (Fig. S19).

Given the overall stability of mapping at the SNP and locus-level, we queried our ability to predict relative phenotypic values across independent populations using allele frequency data alone. Specifically, we computed the relative pigmentation of phenotypic groups (i.e., the light, midpoint, and dark color fractions) both within and across mapping populations iteratively using each independent genotype-phenotype map (see Methods). We found that we predicted relative pigmentation scores with statistically indistinguishable accuracy between, as well as within, mapping populations (Fig 5E; Fig. S23). Strikingly, predicted pigmentation scores maintained their accuracy when excluding variation within the canonical pigmentation genes identified by previous mapping studies, and thus only using variation within loci not previously identified as pigmentation-associated (Fig. S24). This maintenance of out-of-sample prediction accuracy when the canonical genes are removed further demonstrates that the novel trait-loci identified by our mapping are not spurious false positives, but rather harbor pigmentation-relevant variation for the species.

Amidst the strong stability of the genotype-phenotype map across evolutionarily diverged populations (Fig. 5B-D), we still observed several conspicuous instances in which genomic regions harboring pigmentation-relevant variation for one mapping population appeared to lack pigmentation-associated variation in an independent mapping population. Several such instances can be observed in the mirrored Manhattan plot in Fig. 5A, where the start and end points of pigmentation loci (indicated by shaded regions) within one environment exhibit non-overlapping start and endpoints with any of the loci identified in the second environment. We suspect that these environment-specific loci potentially indicate the effects of epistasis altering the marginal effect of pigmentation-associated SNPs across mapping populations. In the Supplement, we show that such instances of environment-specific loci are not solely due to technical noise, but in fact environment-specific marginal effects (Fig. S20-21); the exclusion of these loci from cross-treatment genomic prediction of trait evolution does not alter our inferences (Fig. S22). Still, over and above this detectable set of environment-specific mapped loci, the vast majority of loci are shared across independent mapping populations (e.g., 30 of 43 loci identified in the week 10 outdoor population overlapped with at least one of the 42 loci in the week 10 indoor population), and thus the dominant signal in the data is one of strong stability of mapping.

## Discussion

The central goal of this study was to rigorously evaluate the predictability of adaptation. To do so, we leveraged a model complex trait (abdominal pigmentation), a model species (*Drosophila melanogaster)*, and a highly powered, longitudinal study design in an ecologically-relevant setting. Our data demonstrate that a well-resolved genotype-phenotype map can indeed be used to correctly predict if, and in what direction, a complex trait evolves. However, the specific genetic architecture of adaptation (i.e., which loci show signatures of selection during trait evolution) was largely unpredictable based on mapping data alone. Specifically, as our study populations evolved lighter pigmentation in semi-natural, outdoor mesocosms, we observed a net genome-wide increase in the frequency of light-associated alleles (Fig. 4A). But at the level of individual mapped loci, there were instances in which light-associated alleles behaved neutrally, or were even selected against during this period (Fig. 3D). The key implication of this result is that it shows how genetic redundancy in complex traits may allow populations to adapt, despite underlying constraints ^67–69^. From an applied standpoint, it demonstrates that predicting complex trait evolution hinges upon a well-resolved genotype-phenotype map in which loci of moderate to small effect are characterized, as these may be in fact those that drive evolution of trait means in natural settings.

Our study design and results afforded further insights into the underlying processes giving rise to the unpredictable architecture of adaptation we quantified. Specifically, instances in which light-associated alleles were selected against during the evolution of lighter pigmentation suggest these alleles are pleiotropic, with negative genetic correlations between pigmentation and other fitness-associated traits that came under opposing selection throughout the observation period ^10,18,20^. We indeed anticipated this dynamic to play out in our study system, as a plethora of fitness-relevant traits (e.g. fecundity, developmental rate, and starvation resistance) evolve in concert to, and share underlying genetic correlations with, pigmentation ^43,53,54,70^. Furthermore, the canonical pigmentation locus *bric-a-brac*, for which we identified signatures of selection during trait evolution that were in direct contrast to what was predicted based on our mapping (i.e., a net decrease in frequency of light-associated alleles during lighter pigmentation evolution; Fig. 3D), is a well-documented pleiotropic regulator, with documented effects on traits ranging from pattern formation in limbs to ovary morphogenesis ^61,62,71^. Notably, such pleiotropic constraint may not necessarily be a function of a single allele, but representative of a haplotype-level pleiotropy ^49^ where alleles conferring lighter pigmentation may be in tight linkage to those associated with another fitness-associated trait that is strongly selected against over the same time period.

An additional facet of complex trait variation that may have further exacerbated the unpredictable genetic architecture of adaptation are genotype-by-environment interactions (GxE). Specifically, we conducted trait mapping in common garden conditions (25°C, 12hrL:12hrD) to standardize the environment in which we quantified the genotype-phenotype map across time points and treatments. Trait evolution, however, proceeded in the context of continually shifting abiotic and biotic conditions where the presence of GxE may continually alter the marginal effect of a locus on trait variation (Fig. 1). In other words, the marginal effect of a locus on trait variation in the conditions in which it was mapped may differ from that in the environment in which the trait comes under selection. We indeed suspect this dynamic to, at least in part, explain instances in which light-associated alleles behave neutrally as the population evolves lighter (grey bars in Fig. 3D). However, it is highly unlikely that such GxE induces the behavior we observed at several large-effect loci in which dark-associated alleles increased in frequency while the population became lighter, counter to expectations (blue bars in Fig. 3D). Rather, this ‘counter-gradient’ behavior is a hallmark of pleiotropic constraint driven by negative genetic correlations among fitness-associated traits^20,21,72^ which, again, was expected to occur in our ecologically-realistic, outdoor system, particularly for those loci with known pleiotropic effects, such as *bric-a-brac*^61–64^.

While the impacts of pleiotropy and GxE were pronounced in our genomic predictions of trait evolution, we found epistasis to have a less detrimental effect on the prediction of complex trait variation overall. Specifically, by re-conducting our high-powered mapping in four independent sets of replicate populations, diverged by evolutionary time and environment of selection (Fig. 2B-C), we quantified a striking stability of trait architecture (Fig. 5). We further showed that genomic predictions of relative phenotypic value using genotype-phenotype maps were equally accurate whether applied within or across genetically diverged mapping populations (Fig. 5E). Therefore, at least within the confines of the magnitude and timescale of evolution manipulated here, epistasis (and/or context-dependent dominance shifts) did not inhibit a general transferability of mapping results across populations diverged in evolutionary time and environment of selection. While this finding may be at odds with some studies evaluating the portability of mapping data across diverged populations, it bolsters more recent work demonstrating increased portability under instances in which trait architecture has been quantified in manners that better control for ancestry and demographic effects, allowing better refinement of trait loci and putatively causal variants^25,65,66,73^.

A key additional component of this study was pairing our outdoor mesocosms with indoor mesocosms that were founded with the same outbred population and evolved in the context of increasing population densities, but in the absence of the multivariate abiotic fluctuations that characterized the outdoor habitat. This allowed us to first systematically test whether direct selection imposed by abiotic variables, such as temperature, underpin the predictable patterns of pigmentation evolution in natural populations^43^. Surprisingly, however, we found correlated patterns of lighter pigmentation evolution across the environments, suggesting that lighter pigmentation evolution is at least in part associated with indirect selection on the reproduction-associated traits (e.g., fecundity) that are favored under increasing population densities^49,54^. Despite this shared phenotypic evolution between environments, we still observed differences in underlying genetic architecture of trait adaptation through time. Specifically, we found that pigmentation-associated loci behaved more predictably in the indoor environments, with light-associated variants more likely to increase in frequency as the population itself became lighter. In contrast, in the outdoor environment, we observe stronger counter-gradient behavior, where a handful of dark-associated variants strongly increase in frequency even as the populations evolved to become lighter (Fig. 4). We hypothesize that this pattern is underpinned by the more complex selective regime of the outdoor environment, such that selection acts on a greater number of traits overall which leads to greater pleiotropic constraint on pigmentation-associated loci ^74,75^. From a practical standpoint, this implies that the nuances of adaptation in the wild may be skewed by studying the process in simplified, lab-based environments and bolsters recent calls to study the ecology and evolution of populations in natural contexts ^76^.

Our insights into the predictability of evolution were only made possible by several key aspects of our methodological approach: our generation of haplotype-informed allele frequencies at extremely high coverage, in combination with a large number of biological replicates per mapping treatment and time point, provided the power to produce well-resolved genetic maps (see Fig. S3-4)^51,55^. Furthermore, our longitudinal analysis of trait variation using a single founding population was the only way to exclude confounding variables such as unique patterns of demographic history and linkage when assessing the role of pleiotropy and epistasis on complex trait prediction ^28,30,65^. Crucially, our accurate genomic prediction of trait evolution hinged upon our high-powered genotype-phenotype map that identified both large-effect, canonical pigmentation genes, as well as moderate-effect, previously undescribed, loci. Our study therefore exemplifies the critical roles of study design and power in the rigorous evaluation of fundamental evolutionary dynamics. It is worth noting that while our study focused on the principles of adaptation via standing variation, likely a primary form of evolutionary change in natural populations ^77^, understanding how these principles are impacted by *de novo* mutations is an exciting area of future research.

Looking forward, our results are pertinent to numerous applications of genomic data: predicting phenotype across disparate populations, inferring past selection using time-series and ancient DNA genomic data, or manipulating contemporary variation to shape future evolutionary trajectories (e.g., augmenting population resilience to future global change conditions). The key takeaway from this study is that high-powered mapping data is indispensable and that evolution in a natural setting can give rise to genetic architectures that controlled, lab-based studies alone cannot predict.

## Methods

### Experimental overview

A genetically diverse population of outbred *D. melanogaster* was generated via outcrossing ten male and female individuals from a total of 76 inbred lines, originally collected from Linvilla Orchards, Media, PA ^50^. The outbred population was allowed to recombine and expand for 4 generations in density-controlled conditions, after which 500 males and 500 females from a single age cohort were used to seed each of 10 outdoor and indoor replicate cages. The outdoor mesocosms consisted of ten replicate, 2x2x2 m mesh cages, each harboring a single dwarf peach tree, and was exposed to natural environmental conditions, insect, and microbial communities ^52^. The indoor mesocosms consisted of ten replicate, ∼0.5 m^3^ cages housed in a temperature-controlled laboratory at the University of Pennsylvania. Constant food was supplied to the replicates within each environment: 400 ml of Drosophila media were provided in 900 cm^3^ aluminum loaf pans within each cage three times per week ^49,52^. Monitoring of phenotypic and genomic variation in the replicates began two weeks after cage founding and continued for a period of ten total weeks and seven generations of evolution. Generation time estimates were produced using a degree day model, a class of phenological models that uses accumulated heat exposure to compute a proxy of biological time (here, the number of generations between two time points), which often does not directly scale linearly with Julian day^78^. The degree day model implemented here was specifically calibrated using published *Drosophila melanogaster* developmental rate data^79^.

Population-wide pigmentation variation was quantified in our panel of inbred lines, the recombinant outbred population after 4 generations of expansion and recombination, and the evolving replicate populations in the outdoor and indoor environments at weeks 4 and 10 (20 July and 1 September). To quantify evolved changes in pigmentation, eggs were collected overnight from all replicate cages (indoor/lab and outdoor/field), after which they were transferred to common garden conditions (25°C, 12L:12D), where they were reared for 2 generations. Following density control of F2 eggs (30 +/-5) in replicate vials, adult female abdominal pigmentation (N = 30 individuals per replicate) was scored as the proportion of pigmentation observed on tergites 5, 6, and 7 ^80^. The sum of these scores across the three tergites provided the total pigmentation score for the assayed individual.

We quantified genome-wide allele frequencies throughout evolution weekly in the outdoor environment and at weeks 4 and 10 in the indoor environment. At each collection point, eggs were collected overnight from each cage and then transferred to common garden conditions where larval development took place. 100 random females of the eclosed adult cohort from each replicate cage were then collected and stored separately in ethanol at -20°C for later DNA extraction and sequencing (see below).

At the week four and ten sampling points, we conducted a tail-based mapping of pigmentation variation for each mapping population. Specifically, we first quantified pigmentation variation within each treatment using a random sample of 100 F2 females (3-6 days post-eclosion) and the visual scoring methods described above^80^. Then, for each replicate across both environments, we segregated individuals into one of three trait value groups: (1) those exhibiting pigmentation within the upper (lightest) 15% of the population-wide phenotypic distribution, (2) those exhibiting pigmentation within the lowest (darkest) 15% of the trait distribution and (3) a random sample from those representative of the midpoint (∼35th-65th percentile) of the trait distribution. Seventy-five individuals from each trait value group and replicate were then stored in ethanol at -20°C for later DNA extraction and sequencing. In total, this resulted in 120 samples available for mapping: three trait value groups across ten replicate cages in each of two environments and sampled at two time points.

### Sequencing and allele frequency estimation

Genomic DNA was extracted for both mesocosm time-series and trait mapping samples using NEB Monarch DNA kit. Following sample QC via nanodrop and qubit, whole genome libraries were constructed using Illumina DNA prep kit. Each sample was sequenced to an equal coverage of 8x using Illumina Novaseq 6000. From the resulting sequencing data, we generated haplotype-informed allele frequencies using previously described methods ^51,81^. This method computes allele frequencies at sites previously identified in the genome sequences of each founding inbred strain using an expectation-maximization algorithm that estimates haplotype frequencies in a pooled sample of mapped sequencing reads. Haplotype inference was conducted in window sizes that varied proportionally to the length of un-recombined haplotype blocks expected as a function of the estimated number of generations since original inbred line outbreeding ^81^. Notably, given our empirical coverage of 8x and this haplotype-aware based method of allele frequency computation, our allele frequency estimates are of the accuracy expected given ∼200x coverage and using standard pooled sequencing methodology ^51,52^. After filtering sites to those with an average minor allele frequency ≧ 0.02 in the founder population, and present in at least one evolved sample at a minor allele frequency ≧ 0.01, we retained 1.9 M SNPs for analysis.

### Genome-wide patterns of variation across environments and throughout evolution

We first interrogated our haplotype frequencies derived throughout the experiment to quantify whether there existed skew in the contribution of specific inbred lines to the outbred population used in founding our mesocosm cage replicates. Specifically, we evaluated derived haplotype frequencies within our founding population, as well as the samples collected throughout the progression of evolution in our mesocosm cages. We used Shannon’s evenness, an ecological metric for the distributional uniformity of individuals among species within natural communities, to quantify skew in the contribution of specific founder haplotypes to the derived outbred population (here, haplotypes are analogous to species and observed frequencies to the relative abundance of each haplotype within the outbred population). We found an observed evenness value of 0.97 (maximum value 1) indicating only a 3% reduction in abundance equality, relative to what would be expected under purely uniform haplotype abundance in our outbred population. This evenness is then visualized as the rank order of haplotype frequencies both in our founding, baseline population, as well as the evolved populations sampled throughout the course of the experiment (Fig. S1-2).

We then assessed genome-wide patterns of variation across all samples used in trait mapping using principal component analysis (*prcomp* function, R) and normalized allele frequency data across all 1.9 M SNPs. We projected samples onto the first two principal components and visualized the data by shaping and coloring points according to different underlying variables (e.g., trait value group, treatment) to infer which variables may dominate variance in allele frequencies across samples. We quantified whether samples segregated along the first two principal components in accordance with trait value group or environment using linear regression and ANOVA, respectively.

### Ancestry-informed association analysis of pigmentation variation

Our general approach to mapping pigmentation was to identify SNPs harboring alleles with systematic frequency changes across the light, midpoint, and dark trait value groups. First, however, it was important to consider that due to the finite number of haplotypes from the founding inbred lines used to derive our outbred population, a significant association between a SNP and trait value group could indicate its biological impact on pigmentation variation and/or its inbred-line ancestry. This effect could appear at the level of entire chromosomes due to long range patterns of linkage, as well as more locally around causal loci. We quantified the magnitude of this effect genome-wide by computing the average contribution of haplotypes coming from each inbred line to each trait value group (i.e., the difference in haplotype frequency between the light and dark tails) and quantified the relationship of this value with the pigmentation score of each inbred line using linear regression and Pearson’s correlation coefficient. Lines that significantly contributed to this relationship were determined via an elimination method, where inbred lines were excluded from the regression analysis in order of magnitude of color fraction enrichment. Any lines eliminated prior to the p-value of the regression coefficient dipping below 0.05 were considered to be significantly associated with color fraction differences.

We interrogated the effect of linkage on SNP mapping further by running two generalized linear models (GLM; logistic link function and quasibinomial error variance). The first model only took into consideration the impact of trait value group on variation in allele frequencies (model formula: allele frequency ∼ trait value; where allele frequencies are normalized by the SNP’s effective coverage and the total number of chromosomes sampled, and trait value is either 0, 1, or 2, where 0 is dark, 1 is midpoint, and 2 is light). If the effects of ancestry on SNP association were pervasive, SNPs identified via this approach would be enriched in those SNPs expected to have elevated light versus dark frequency differences based on their association with particular haplotypes alone. To test this, we selected the top 1% of the SNPs (based on nominal p-value from the GLM) and calculated their expected frequency difference between light and dark color fractions based on estimates of the average contribution of each inbred line to that sample and the association between that SNP and inbred line. We then compared these results to a separate GLM model in which the haplotypic ancestry was accounted for (model formula: allele frequency ∼ trait value + ancestry-predicted frequency). This model then regressed allele frequencies both on pigmentation trait value, as well as each SNP’s expected frequency in the sample, calculated using average genome-wide haplotype frequencies across the whole sample and the presence or absence of the SNP’s alternative allele on each haplotype. In accordance with the previous ancestry-agnostic model, we used a logistic link function and quasibinomial error variance, and allele frequencies were weighted to account for variation in coverage and the finite number of chromosomes sampled. We then compared the density distributions of expected light-dark frequency differences for SNPs identified via each of these distinct models using a KS test.

Following GLM model structure selection, we used permutations to define a final set of trait-associated SNPs. Specifically, we randomly shuffled trait value data and re-ran our regressions N = 100 total times to generate an empirical null distribution. We then considered a SNP significant if its nominal p-value from the empirical GLM fell in the upper 99.95^th^ percentile of the permuted p-value distribution (corresponding to an empirical FDR threshold < 0.0005).

The set of pigmentation-associated SNPs identified via GLM was then used to quantify the number and distribution of unlinked loci underpinning trait variation. We developed an iterative procedure for this, whereby we first identified the most significant SNP on each chromosomal arm, as well as its surrounding, locally un-recombined region (defined by the recombination rate and number of generations since cage founding^51^). We then eliminated all trait-associated SNPs on the chromosomal arm (i.e., those with a significant GLM p-value) whose relative presence in the light and dark fractions can be explained by haplotype frequencies (i.e., ancestry) in the locally un-recombined focal region alone. As the significance of these SNPs could be driven either by linkage to the focal top SNP or by variation in the trait itself, we eliminated these and defined the trait-associated locus as the locally unrecombined region (as defined using number of generations since founder line outbreeding and genome-wide recombination rates) around the lowest p-value SNP. The specific threshold used in SNP elimination was the top 10% most “light” predicted SNPs and the top 10% most “dark” predicted SNPs. This threshold was defined via permutations: we iteratively subset the 10 replicates into two equal subsets (N = 5). In one subset, termed ‘real-data’, light-dark haplotype frequency differences within the focal region were computed as above. In the other subset, the dark-light haplotype frequency difference was calculated (i.e., termed ‘sign-flipped’ data). In each subset we sequentially eliminated percentiles of SNPs starting with the top and bottom 2.5% of expected light-dark frequency differences and selected our threshold as the percentile at which the number of true significant SNPs eliminated in the real data matched that of the sign-flipped data (Fig. S25). From the set of remaining SNPs (those not eliminated in the first iteration), we identified the SNP with the most significant nominal p-value and repeated the procedure. This was iterated until we exhausted all trait-associated SNPs on the chromosome. For each chromosomal arm, we then quantified how the number of trait-associated SNPs decreased as a function of these iterations (Fig. S26). Finally, we merged all loci with overlapping breakpoints to procure a final set of unlinked and independent trait-associated loci. A visualization of this clustering procedure is presented as Supplementary Figure 10.

We additionally explored which candidate genes may be underlying the identified loci. Specifically, we identified genes within each locus with a GO term associated with pigmentation (excluding eye pigmentation) or cuticle development/molting. For each of the identified genes, we then quantified whether the number of pigmentation-associated SNPs within the gene (identified via mapping) was greater than expected based on their genome-wide abundance using a binomial test with a Benjamini-Yekutieli false discovery rate correction.

### Power simulations of highly-replicated tail-based mapping

We evaluated the power of our tail-based mapping using a series of simulations in the forward-in-time population genetic simulator, SLiM^82^. We specifically aimed to assess the power of our study design to detect loci of a range of empirically-informed effect sizes, and how our study’s power specifically compares to more standard, individual-level GWAS approaches (e.g., using lines available from the Drosophila Genetics Reference Panel). The universal variables established and held constant across simulation were as follows: 10 quantitative trait loci spread across a 20 Mb chromosome, each of which was separated by at least 1 Mb from adjacent loci. The loci were assigned a range of effect sizes, which in SLiM are encoded as the percentage of variance explained (PVE), to determine the effect of an allele on the complex phenotype. We evaluated PVEs ranging from 0.0001 to 0.05, which represented the range of detection limits (from undetectable to detected in 100% of simulation scenarios) for both the tail-based and individual-level GWAS approaches. Our 10 modelled loci account for ∼10% of the genetic variation, so the remaining ∼90% of genetic variation and its contribution to a fly’s phenotypic value is modelled as a simple random variable for each individual. The environmental variance and genetic variance encoded into the model simulations were determined empirically and based on quantification of pigmentation in the focal inbred line panel used in this study: the environmental variance is the square of the average standard deviation of pigmentation within individual inbred lines (SD ≈ 1.24); the genetic variance is the square of the standard deviation of pigmentation across all our inbred lines (SD ≈ 5.07), quantifying the variation due to the existing genetic differences between the founding lines. From these values, we computed the broad-sense heritability of pigmentation to be ∼ 0.89.

We then determined our power to identify the simulated trait loci using a replicated, tail-based approach as conducted in this experiment, as well as an individual-level GWAS using inbred lines, a common approach to complex trait mapping in *D. melanogaster*. Simulations of the tail-based approach were carried out identically to our experimental methodology: we initiated simulations with 76 haplotypes each randomly assigned a presence/absence of the causal allele at each of the 10 trait-associated loci (the number of lines possessing each causal allele ranged across simulations to quantify the impact of causal allele frequency on locus identification). Twenty individuals from each line were then mixed and allowed to reproduce and expand for four generations to a population size of 20,000. At this stage, sets of 1000 individuals were randomly selected to serve as the founding population for a set of replicate populations, which were then each allowed to evolve and expand to a size of 50,000 over the course of eight additional generations under a Wright-Fisher model of evolution. The population sizes were determined empirically from census counts throughout the experiment. Following this burn-in period, 1500 individuals were randomly selected per replicate population, their pigmentation computed (by summing the number of causal loci each individual harbored and their relative effect sizes, and while accounting for the environmental and genetic noise), and then segregated into the 15% darkest, the 15% lightest, and midpoint of the trait value distribution. As with our experimental design, seventy-five individuals per trait value group and replicate were randomly selected and used to compute the allele frequencies. These data were then used to map the association between each SNP and trait value group using a generalized linear model with quasibinomial error variance structure. These simulations were conducted over a range of replicate population numbers (from 1 to 10), allowing us to develop an intuition regarding how the power of this approach scales with biological replication, as well as over a range of minor allele frequencies at causal loci, ranging from rare to common (0.02, 0.05, 0.1, 0.2, and 0.3).

We next evaluated the detection power of the same set of simulated loci using the more common approach of individual-level, inbred line GWAS. As above, each inbred line simulated for the GWAS was assigned the presence/absence of each causal allele, and their pigmentation determined based on the sum of the presence of causal alleles and their relative effect sizes. Twenty individuals were phenotyped per GWAS line, and the mean of these individuals’ pigmentation scores was used as the phenotypic value for the inbred line. This level of replication was chosen to mirror the replication in Dembeck *et al*. (2015), in which the authors phenotyped 10 females per line ^49^. To be conservative with our comparison, we doubled the number of individuals phenotyped. Mapping was then conducted using a simple linear model regressing genotype (0 for non-carriers of causal allele, and 2 for carriers) on phenotype. We evaluated GWAS mapping power over a range of sample sizes (from 200 to 1000 inbred lines) and, as above, over a range of minor allele frequencies at causal loci (0.02, 0.05, 0.1, 0.2, and 0.3).

### Comparison of mapping to previous association studies

We leveraged results reported by two previous genome-wide association studies ^47,48^ to quantify the consistency of mapping of pigmentation across disparate populations and studies. In brief, Bastide et al. (2013) conducted a pooled-GWAS approach using wild populations of *D. melanogaster* from Austria and Italy, and focused on female pigmentation on tergite 7. Dembeck et al. (2015) conducted a GWAS using individual *D. melanogaster* lines from the Drosophila Genetics Reference panel ^83^, focusing on pigmentation variation of tergites A5 and A6. We evaluated whether all SNPs identified as trait-associated in the Dembeck study and the top 100 SNPs from Bastide et al. exhibited more overlap with our unlinked loci than expected by chance. Specifically, we compared the number of SNPs falling within each locus to a random set of SNPs within the chromosome of the same test set size as those obtained from the comparison studies. We iterated this process 1000 times and generated an empirical p-value given the observed overlap, relative to the randomly generated 1000 overlaps. We then re-conducted this analysis, but excluded those within loci containing *tan*, *ebony*, and *bab* to infer the extent to which shared signal across studies was driven by these large effect genes.

### Predicting evolution using mapping results

We assessed the predictability of the architecture of adaptation by quantifying whether mapped SNPs associated with lighter pigmentation increased in frequency as the experimental populations evolved to become lighter. Specifically, we computed the frequency shifts for all light-associated alleles at pigmentation-associated SNPs between weeks 3 and 10 of evolution in the outdoor mesocosm separately for each of the unlinked, trait-associated loci. We compared the resulting distribution to one derived via a set of matched control SNPs (matched on chromosome and starting frequency (± 2.5%), and excluding those SNPs within 500 Kb of the respective target SNP) using a binomial test. The p-values from these tests were corrected for multiple hypothesis testing via Benjamini-Yekutieli correction. In effect, we quantified whether the proportion of light-associated SNPs within each locus shifting in a manner concordant with that predicted by mapping (i.e., increasing in frequency as lighter pigmentation evolved) was significantly greater than expected based on our matched control distribution.

We then compared how our genomic predictions of trait evolution differed between the semi-natural outdoor mesocosms and relatively simple indoor environment. First, we quantified the trajectory of trait-associated SNPs (identified in our outdoor, week 10 mapping population) separately in the outdoor and indoor mesocosms throughout the period of lighter pigmentation evolution (week 3 through 10). We compared each resulting distribution to a set of matched control SNPs using a binomial test to infer whether the direction and magnitude of light-favored allele frequency shifts exceeded background allele frequency movement. This analysis was then re-conducted separately for SNPs within each of the top loci on chromosomal arms 2L and 2R, as well as the loci containing the genes *bab*, *ebony*, and *tan* on chromosomal arms 3L, 3R, and X, respectively. Furthermore, the distribution of trait-associated SNP trajectories throughout trait evolution within each environment were compared directly to each other using a pairwise t-test.

Finally, we generated genomic predictions of trait evolution using an approach that explicitly accounted for the observed effect size of each SNP on trait variation observed via mapping. Specifically, we summed the frequency shifts of light-associated alleles, this time weighting each allele’s shift by the associated SNP’s GLM regression coefficient. This summation then represents a genomic prediction of relative phenotypic shift in population pigmentation based on our genotype-phenotype map, whereby positive values indicate the evolution of lighter pigmentation and negative values, darker pigmentation. We bootstrapped these estimates N = 1000 times, and their significance was then assessed using Z-scores computed via a comparison of the estimate to a distribution generated via N = 1000 matched control SNP sets (matched on chromosome and starting frequency) that did not display evidence of a significant pigmentation association in our mapping. We considered the empirical, bootstrapped value significant if its confidence intervals did not overlap with the matched control distribution.

### Stability of mapping throughout evolution and across environments of selection

We determined the stability of the genotype-phenotype map across independent mapping populations (i.e., each mapping environment and timepoint) using several approaches. First, we re-conducted our mapping procedure, as described above, for the indoor weeks 4 and 10, as well as the outdoor week 4 mapping population. Next, we asked if the proximity of loci identified across mapping populations were more similar than expected by chance. To address this, for each of the four genotype-phenotype maps we identified the top SNP of each locus (the SNP with the lowest p-value, and also the locus midpoint) and summed the number of times it fell within a locus identified in one of the independent mapping populations. This then produced a score, ranging from 0 to 3, for each locus and these scores were summed across all loci to produce an empirical overlap score for each mapping population (note that to increase power of this analysis, we selected focal SNPs using the set of pre-merged loci for each mapping population). To determine if the observed overlaps were greater than expected by chance, we generated an empirical null distribution via random sampling of sets of SNPs for each genotype-phenotype map and re-computing overlap scores with each independent mapping (a procedure iterated N = 2000 times for each of the four genotype-phenotype maps). We then interrogated whether the distance between locus-defining midpoint SNPs across sets of loci were closer than expected by chance. Specifically, as before, we first identified the top SNPs for each set of loci from every genotype-phenotype map, and then computed its distance to the closest top SNP of each of the remaining three mappings. This distance metric was averaged to produce an average proximity score which, as above, was compared to a distribution of scores derived via a random set of SNPs (N = 2000 times).

We next aimed to quantify the extent to which the identity of specific pigmentation-associated SNPs was shared across independent mapping populations. To do so, we compared the overlap in identified SNPs within and between independent mapping populations. To conduct this analysis in a balanced manner (i.e., compare sets of SNPs identified using the same number of replicate cages/populations), we iteratively subset our set of 10 replicates from each mapping population into groups of 5 replicates and identified pigmentation associated SNPs in each subset using the ancestry-informed regression introduced above. This was iterated with each possible combination of subset replicates, yielding 126 total sets of significant SNPs per mapping population. The percentage of shared SNPs was then calculated for subsets of the same mapping population (e.g. overlap between pigmentation-associated SNPs identified using different subsets of 5 replicate populations from the outdoor, week 10 replicates), and compared to those observed between independent mapping populations (e.g. overlap between sets of pigmentation associated SNPs identified using 5 replicate populations from outdoor week 10 replicates and sets of SNPs from independent treatment and/or time point). Whether the amount of overlap in identified SNPs within vs. between mapping populations differed was evaluated with a planned contrast using a one-way ANOVA model.

We next sought to quantify whether there existed mapped loci with environment-specific marginal effects on pigmentation variation. Accordingly, we first isolated those loci from the outdoor and indoor week 10 mapping populations that had non-overlapping breakpoints with any of the loci in the opposing environments (N = 13 total unique outdoor loci, and N = 13 total unique indoor loci). We then identified all pigmentation-associated SNPs within each unique locus and quantified the light-dark frequency difference for the light-associated allele for the environment in which it was identified, as well as the environment that did not contain the locus. Evidence for environment-specific marginal effects of each locus was then determined by comparing the distribution of light-dark frequency shifts for each locus between environments (using a Wilcoxon Signed Rank Test), as well as to a distribution derived from an equally sized set of matched control SNPs (using a Mann Whitney U test).

Finally, we sought to determine if genotype-phenotype maps produced at a particular point in evolution, and within a particular environment of selection, could correctly predict phenotypes of individuals sampled at an independent time and environment. To do so, we used those SNPs identified in our outdoor week 10 mapping and calculated the relative expected phenotypic score for trait value groups (e.g., light, midpoint, and dark color fractions) within this focal mapping, as well as all remaining, independent mapping populations. Specifically, for each light-associated allele at all pigmentation-associated SNPs, we calculated the difference in frequency between the baseline population and each pooled sample. This allele frequency difference was weighted in accordance with the SNP’s effect size on trait variation as computed via mapping (i.e., by multiplying the light-favored allele frequency change by the regression coefficient from the ancestry-informed GLM). These values were then summed across all SNPs for each sample, providing a metric of the relative lightness of the sample, as predicted by mapping (values greater than 0 indicating lighter pigmentation and less than 0 darker). We then bootstrapped this value 1000 times for each sample. The accuracy of relative predicted phenotypic scores was then evaluated using three criteria: (i) whether the dark samples had, on average, predicted phenotypic values below 0, (ii) whether the light samples had, on average, predicted phenotypic values above 0, and (iii) if there existed a significant linear association between trait value group and relative predicted phenotypic value. The latter was tested using a simple linear regression with the model trait value group ∼ phenotype prediction, where trait value groups were assigned a numeric value (dark = 0, midpoint = 1, and light = 2).

#### Code and data availability

All code necessary to replicate the analyses presented in this manuscript are available at the following: https://github.com/jrhodes613/PigMap2021.

Founder line sequences are available at the NCBI with the accession ID PRJNA722305. Upon manuscript acceptance, sequences generated via pooled sequencing for this study will be available at the NCBI with the accession ID PRJNA1031645 and phenotypic and allele frequency data will be uploaded to a dryad repository.

## Author contributions

Conceptualization: MCB, DAP, PS

Experimental methodology: MCB, SB, JB, DAP, PS

Formal analysis: MCB, JASR

Writing of manuscript: MCB, JASR, DAP, PS

Manuscript editing: all authors

## Acknowledgements

We thank Lauren McIntyre, Jonathan Pritchard, and all members of the Petrov and Schmidt lab for helpful feedback of experimental design and analysis and interpretation. We thank Hayes Oken for experimental support, Marianna Karageorgi and Emily Behrman for assistance with the degree day model, as well as Caitlynn T. Tran and Andy V. Huynh for assistance in pigmentation phenotyping. We thank Heloise Bastide for an extended list of pigmentation-associated loci and results from Bastide et al 2013 (doi: 10.1371/journal.pgen.1003534).

## Supplementary Information for Smiley-Rhodes* and Bitter*, *et al*

### Supplementary Figures

**Supplementary Figure 1.**
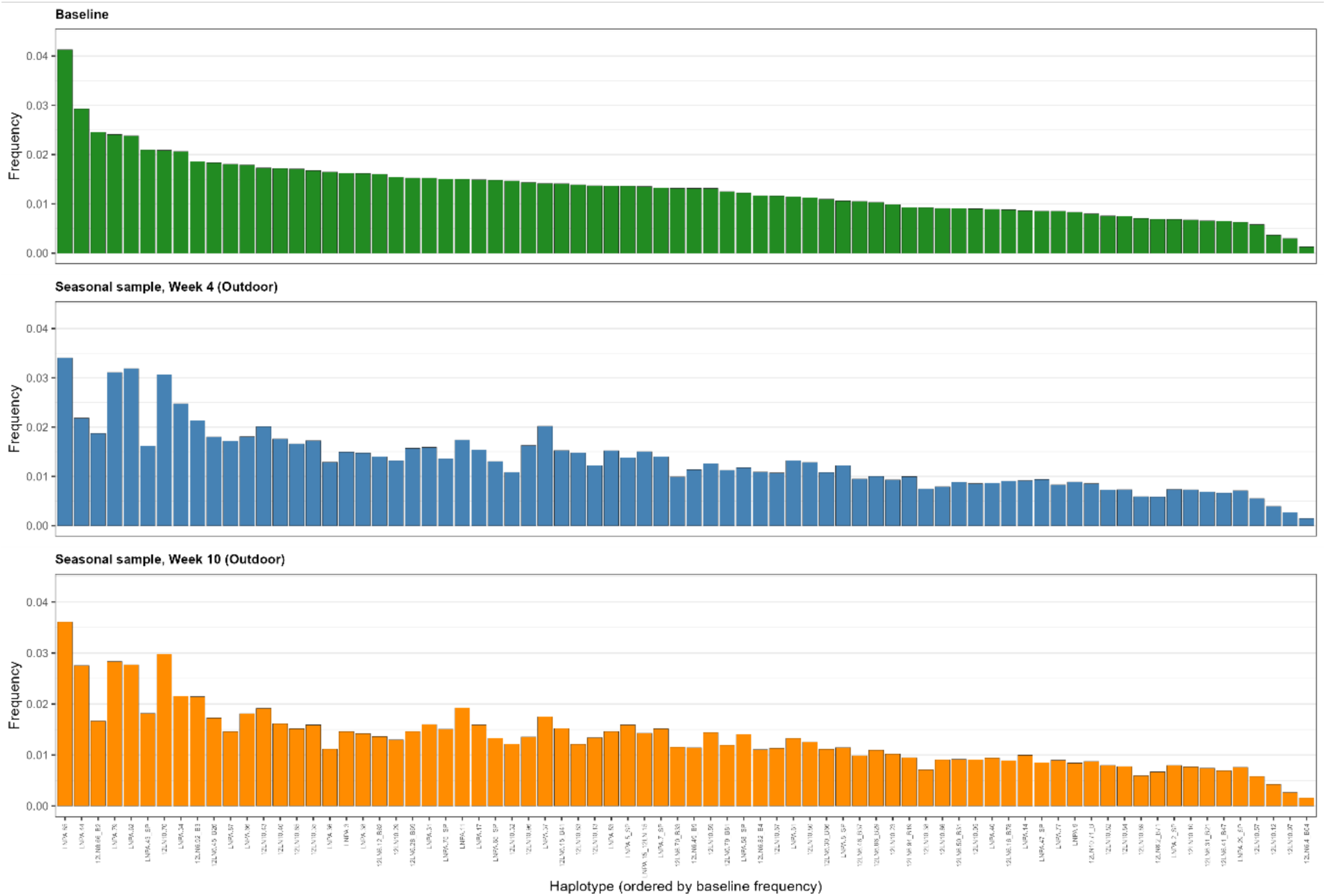
Inbred-line haplotype frequencies across baseline and evolved outdoor mesocosm populations. Distribution of inbred line haplotype frequencies samples taken from the baseline outbred population before cage seeding (green) and evolved outdoor mesocosm populations collected at weeks 4 (blue) and 10 (orange) of the experiment. Inbred lines are ordered from most frequent to least frequent in the baseline population.

**Supplementary Figure 2.**
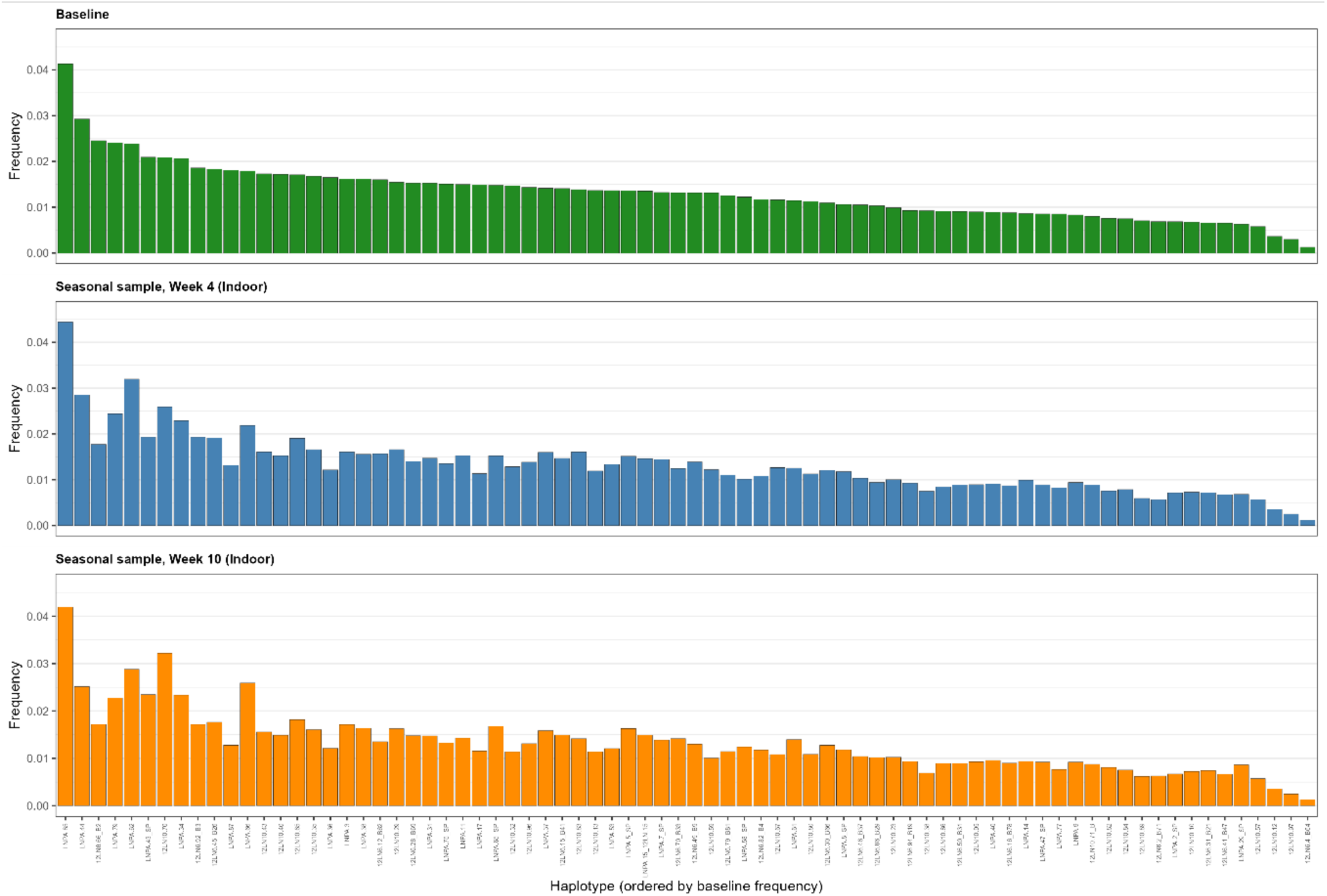
Inbred-line haplotype frequencies across baseline and evolved indoor mesocosm populations. Distribution of inbred line haplotype frequencies samples taken from the baseline outbred population before cage seeding (green) and evolved indoor mesocosm populations sampled at weeks 4 (blue) and 10 (orange) of the experiment. Inbred lines are ordered from most frequent to least frequent in the baseline population.

**Supplementary Figure 3.**
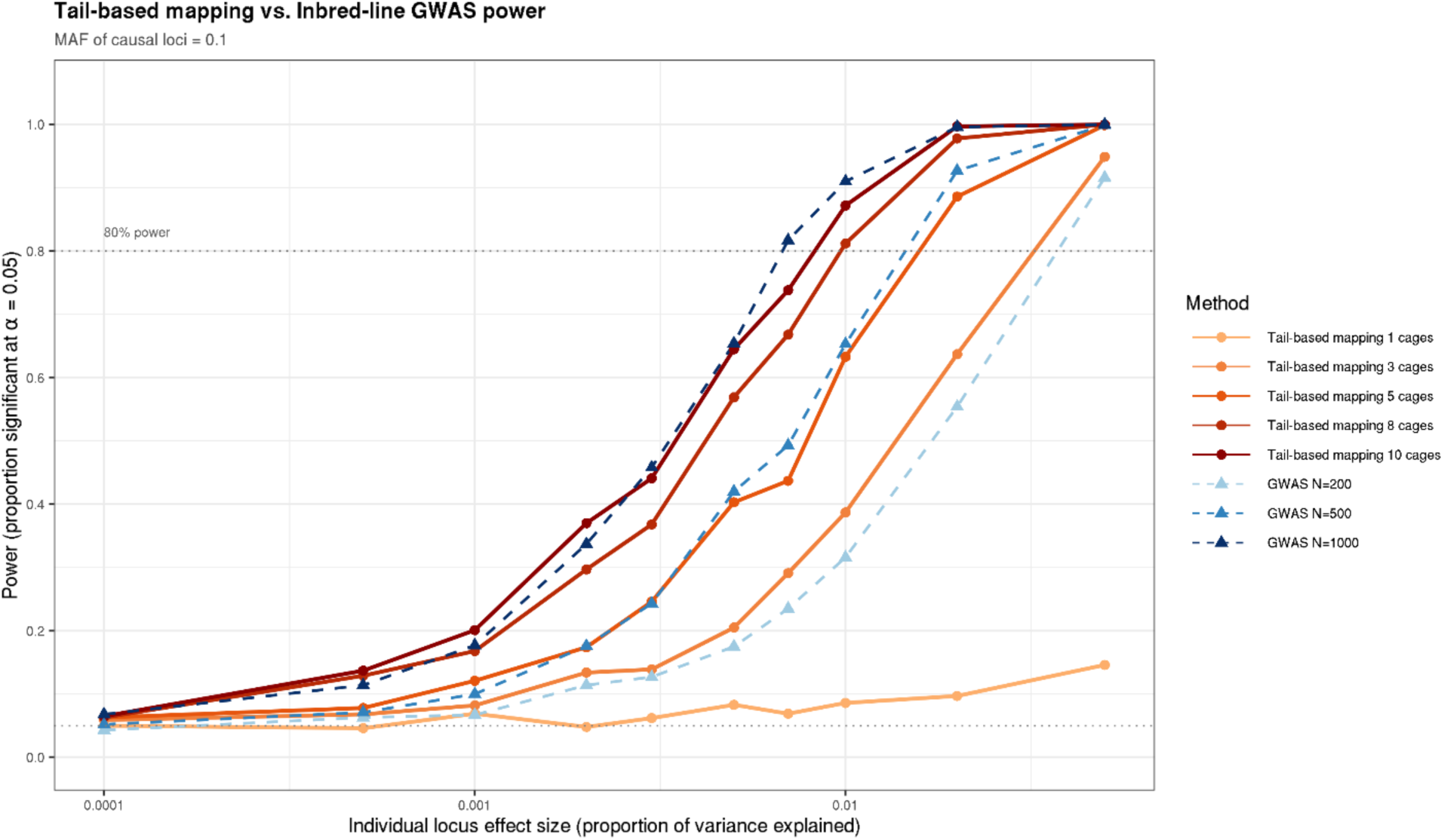
Power simulations of tail-based mapping: impact of biological replication. Detection power for 10 simulated trait-associated loci, ranging from an effect size (proportion of trait variance explained) of 0.0001 to 0.05, under a range of replication scenarios for tail-based mapping (N = 1-10 independent mapping populations) and standard, individual-level GWAS using inbred lines. Here, power is quantified as the proportion of 1000 replicate simulations in which each causal locus was correctly inferred via regression-based association analysis (see Methods). For all simulations presented here, causal allele frequencies were encoded to 0.1 prior to initiation of evolutionary simulation. Simulation results from tail-based mapping design are in warm colors with solid lines, and simulations of results from standard, inbred line-based GWAS are in blue with dotted lines.

**Supplementary Figure 4.**
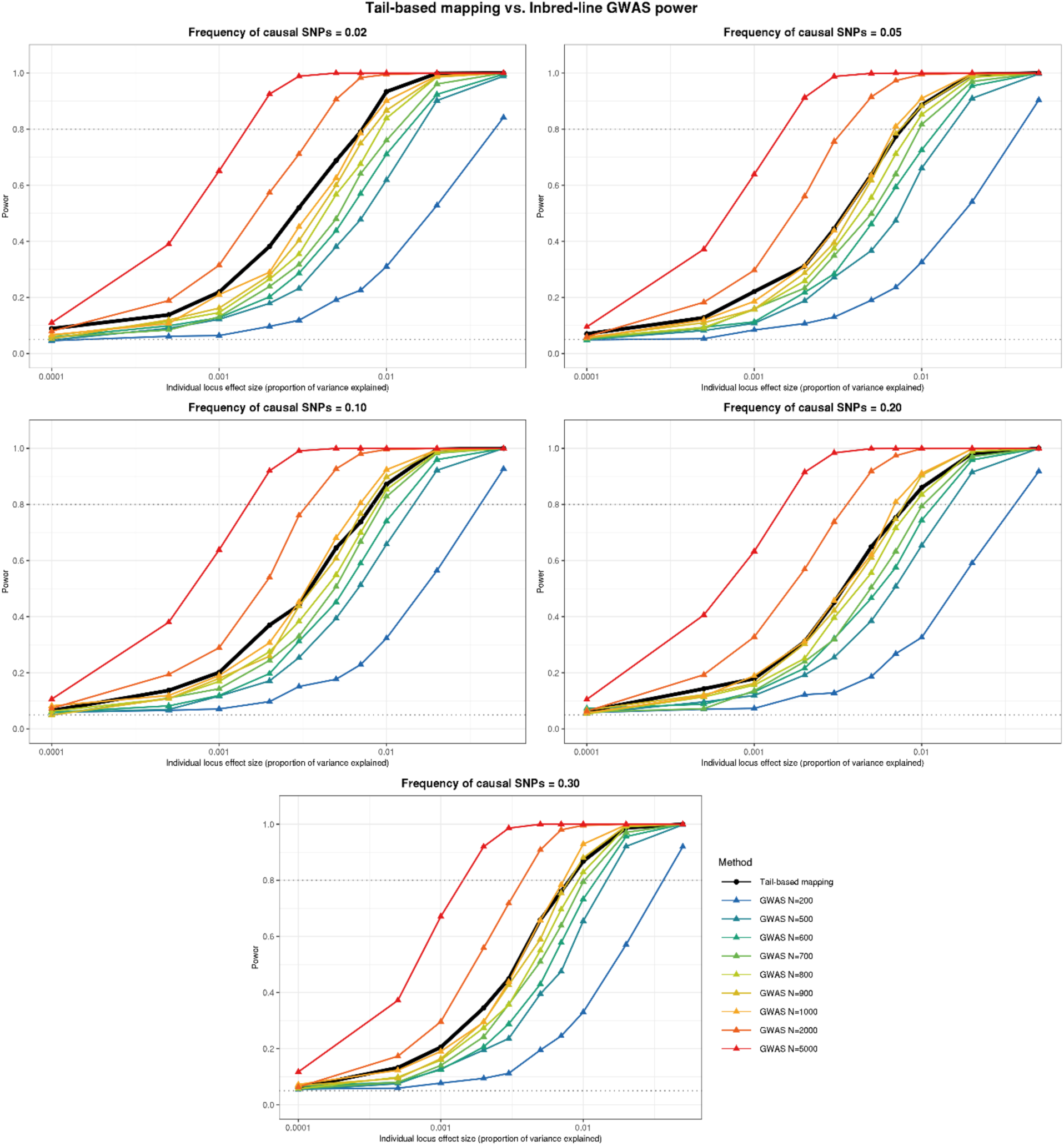
Power simulations of tail-based mapping: impact of causal allele frequency. Detection power for 10 simulated trait-associated loci, ranging from an effect size (proportion of trait variance explained) of 0.0001 to 0.05, under a range of replication scenarios for standard, individual-level GWAS using inbred lines (N = 200-5000 independent inbred lines) and our tail-based mapping approach (N = 10 replicate cages). Here, power is quantified as the proportion of 1000 replicate simulations in which each causal locus was correctly inferred via regression-based association analysis (see Methods). Panels correspond to simulations run under different starting frequencies of causal alleles (from 2-30%). The simulation corresponding to the experimental approach of this study (tail-based mapping, with 10 replicate populations) is presented in black, while individual-level GWAS results are colored according to the number of inbred lines simulated and used in mapping.

**Supplementary Figure 5.**
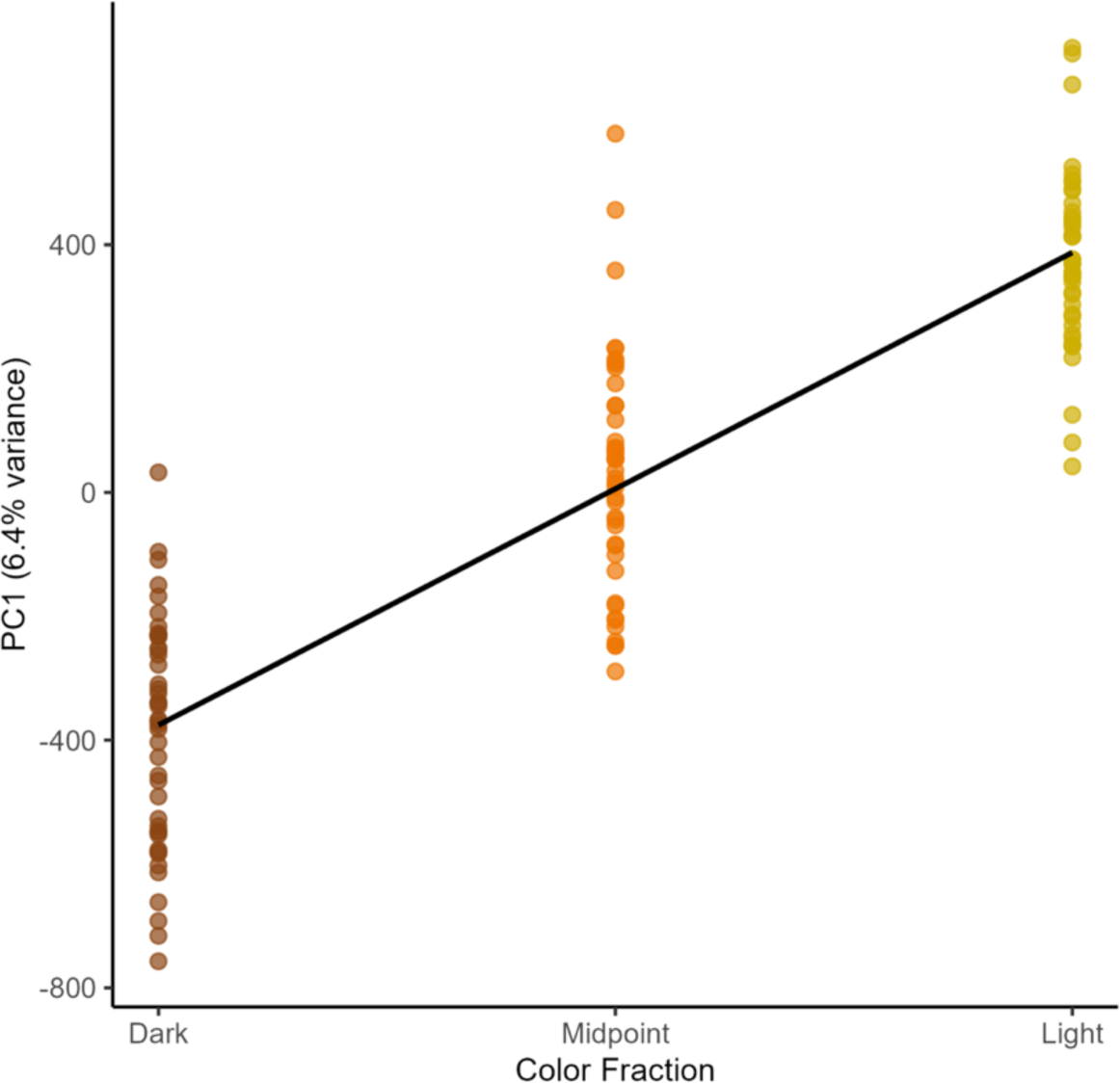
Relationship between color fraction and PC1. Samples plotted by color fraction (dark, midpoint, or light) and the coordinate along the first principal component generated via PCA of allele frequency data across all SNPs and samples used in mapping. A linear regression line (line in figure) between trait value and PC1 is significant (R^2^ = 0.762; p << 0.001).

**Supplementary Figure 6.**
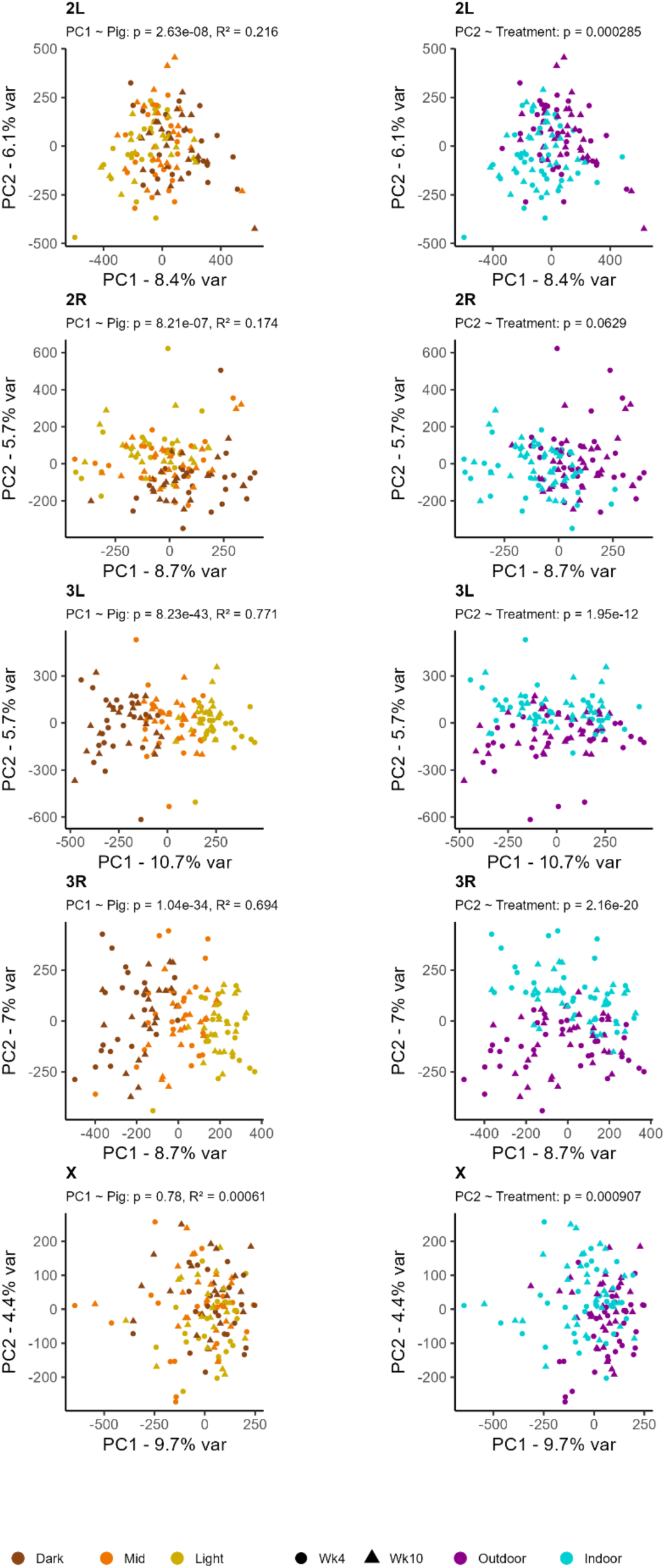
PCA by chromosome. Projection of samples used in trait mapping onto the first two principal components generated via PCA of allele frequency data from all SNPs located on each chromosomal arm. In the column on the left, samples are colored by the color fraction from which they originate (dark, midpoint, or light) and the shape of the points corresponds to the timepoint when the sample was collected. In the right column, samples are colored by treatment.

**Supplementary Figure 7.**
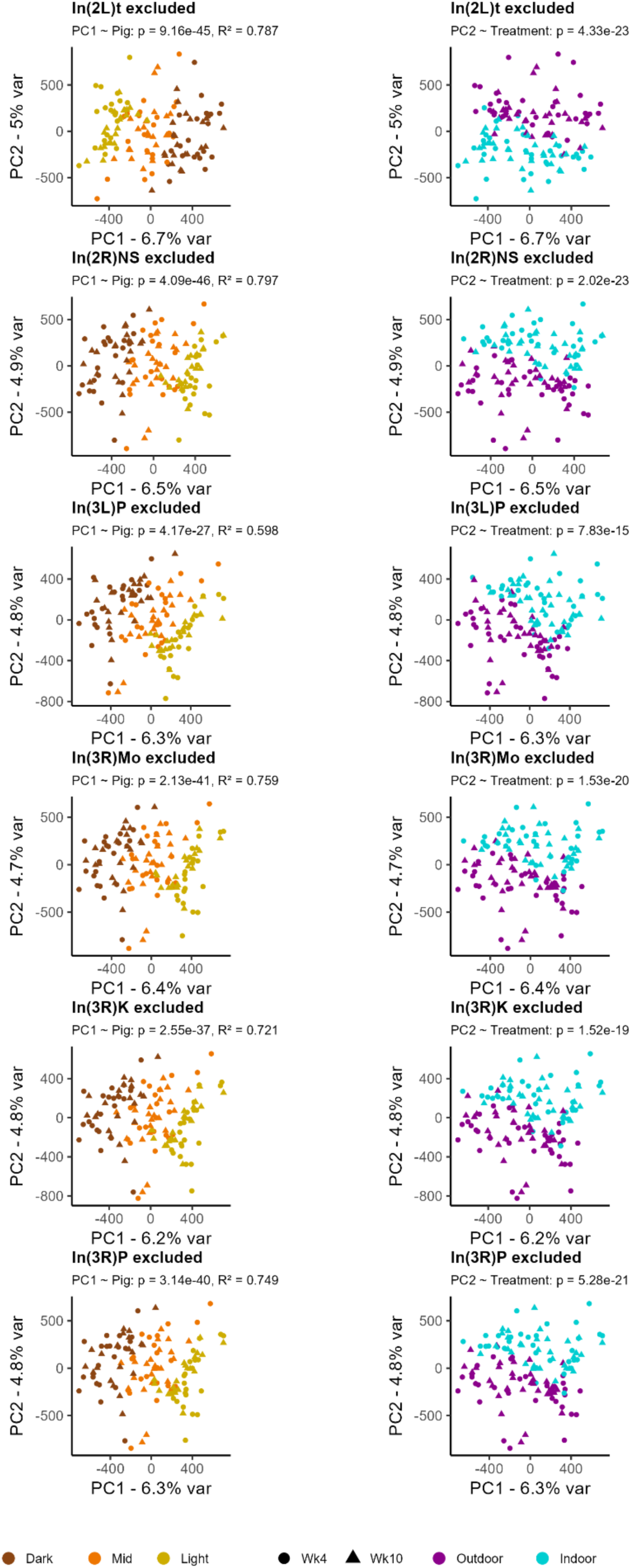
PCA without variation within inversions. Projection of samples used in trait mapping onto the first two principal components generated via PCA of allele frequency data from all SNPs except those located in each of six common inversions found in wild populations. In the column on the left, samples are colored by the color fraction from which they originate (dark, midpoint, or light) and the shape of the points corresponds to the timepoint when the sample was collected. In the right column, samples are colored by treatment.

**Supplementary Figure 8.**
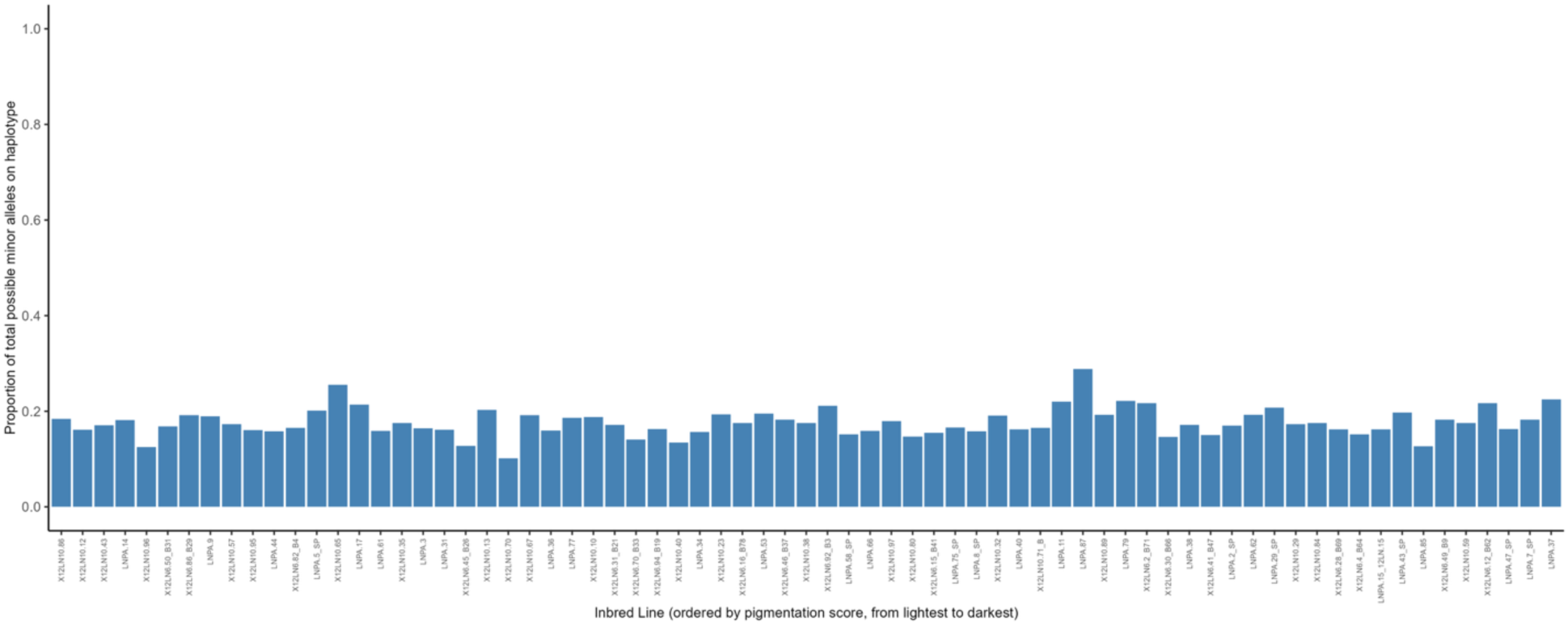
Distribution of pigmentation-associated SNPs across inbred reference panel. The proportion of all pigmentation-associated, significant SNPs (minor allele, identified via our ancestry-informed GLM conducted on the week 10, outdoor mapping population) present on each of the 76 founding inbred lines. Inbred lines are ordered from lightest to darkest pigmentation, as quantified via manual scoring. As can be observed, those SNPs identified by our ancestry-informed GLM are not exclusive to either the lightest or darkest lines, but rather present at appreciable frequency across all haplotypes.

**Supplementary Figure 9.**
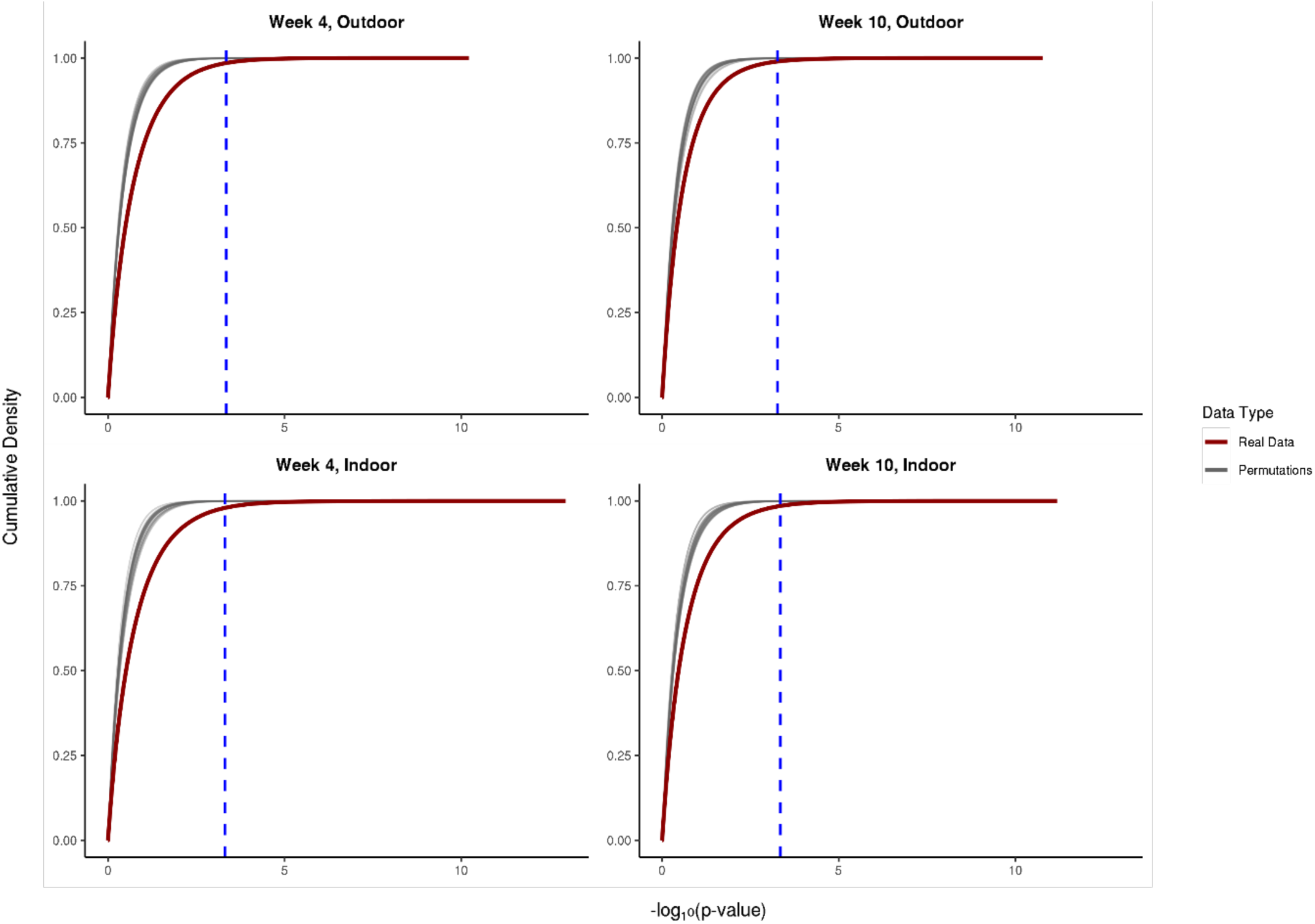
GLM empirical and permuted p-value cumulative distribution functions. CDFs of log10 p-values of SNPs resulting from a GLM performed with real data (red) and 100 GLMs performed with shuffled, permuted data (grey). The cutoff, determined as the top 0.05% of permuted p-values, is drawn as a blue dotted line for each timepoint and treatment.

**Supplementary Figure 10.**
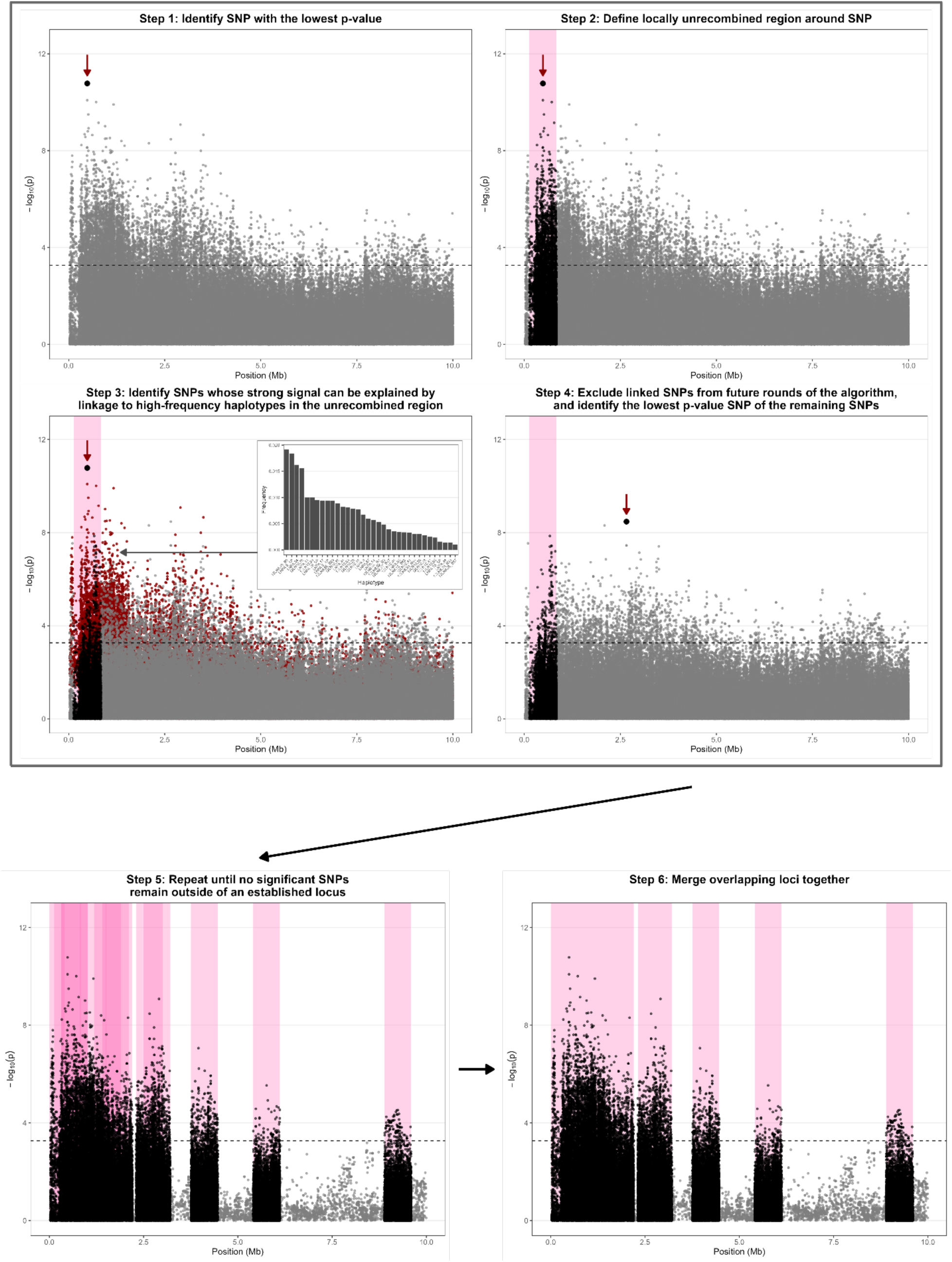
Diagram of algorithm for determining independent, unlinked loci. The establishment of unlinked, pigmentation-associated loci is conducted independently on each chromosomal arm. We start our clustering algorithm by identifying the lowest p-value SNP (derived from our ancestry-informed, generalized linear model) (Step 1). We then define the locally unrecombined region around the focal SNP, given the number of generations of recombination since the initiation of inbred line outcrossing, and the species’ recombination rate (Step 2). Next, we eliminate potentially spurious association signal, chromosome-wide, due to linkage with the focal region defined in Step 2. This is achieved by identifying and eliminating SNPs with a frequency difference between the light and dark trait value groups that is predictable based on their association with the haplotypes driving light vs. dark differentiation within the focal region (Step 3). Thus, after this first round, we are left with our first trait-associated locus (the locally unrecombined region around the lowest p-value SNP), and all SNPs with signature of trait association that may be due to linkage with this locus have been removed. We then proceed to identify the remaining SNP with the lowest p-value (Step 4), and then repeat Steps 2-4 until no SNPs passing our significance threshold (dashed line, determined via empirical permutations) remain (Step 5). Finally, loci with overlapping start/end points are merged into a single locus (Step 6).

**Supplementary Figure 11.**
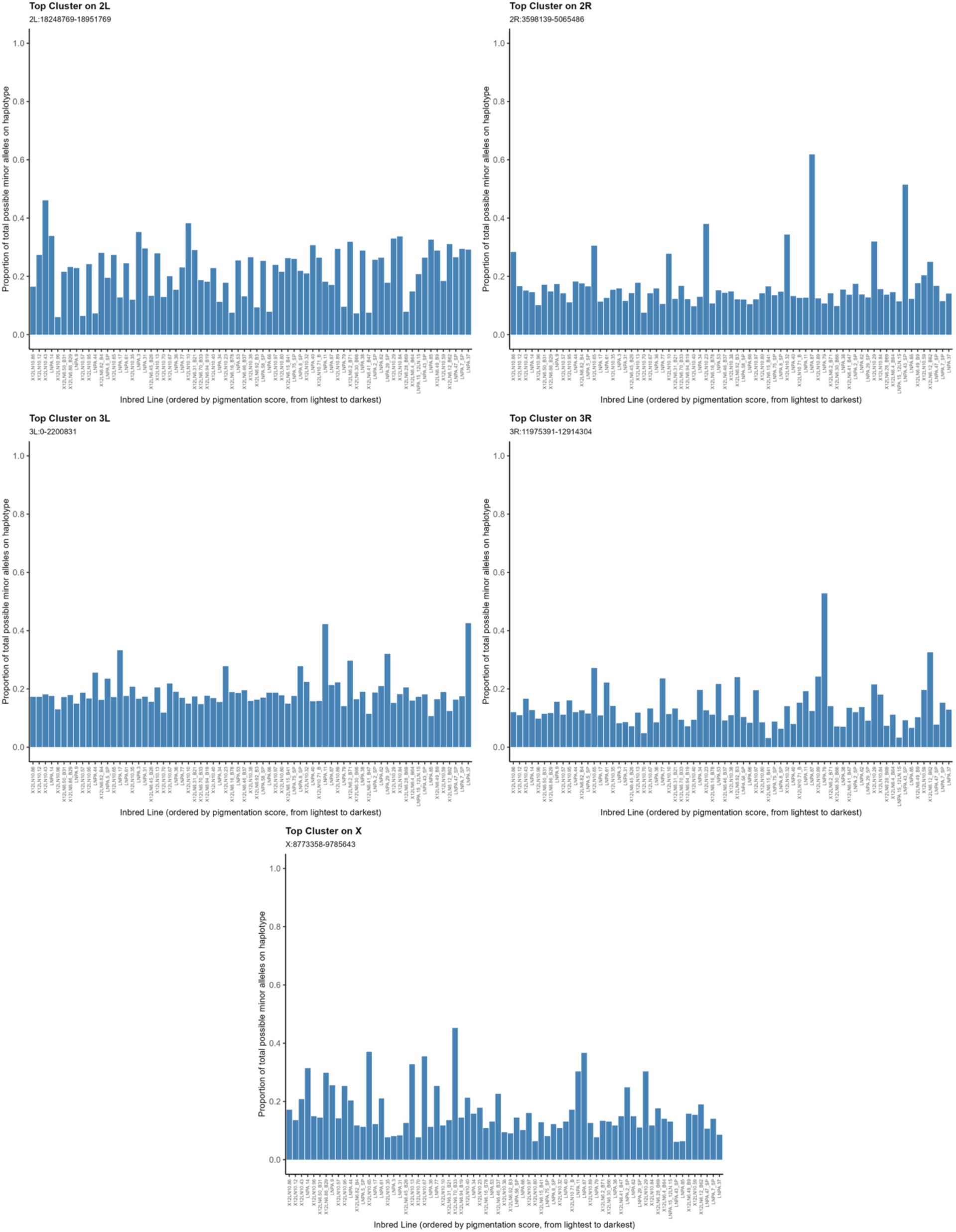
Distribution of pigmentation-associated SNPs across inbred reference panel for the largest-effect, independent loci identified on each chromosomal arm. The proportion of all pigmentation-associated SNPs within each of the top independent loci (clusters) on each chromosomal arm (minor allele, identified via our ancestry-informed GLM conducted on the week 10, outdoor mapping population) present on each of the 76 founding inbred lines. Inbred lines are ordered from lightest to darkest pigmentation score as quantified via manual scoring.

**Supplementary Figure 12.**
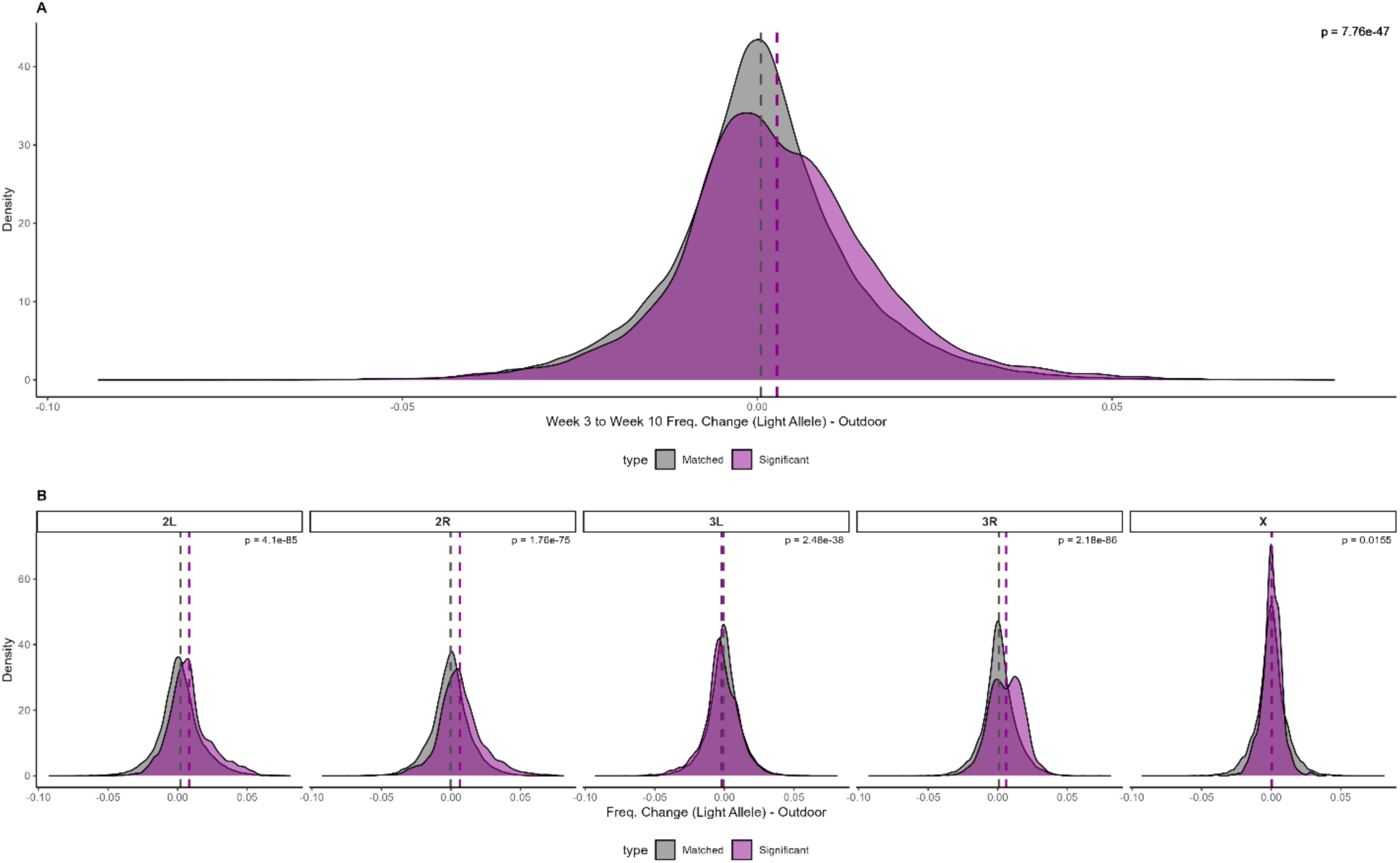
Seasonal shifts of trait-associated SNPs in outdoor cages. A) Distribution of light-associated allele frequency shifts at all trait-associated SNPs (identified at Week 10 in outdoor cages) between Weeks 3 and 10 of evolution in the outdoor field environment. Distributions of frequency shifts at trait-associated SNPs are dark magenta, while distributions derived from matched controls are grey. Reported p-values are derived from a comparison of trait-associated and matched control SNPs using a binomial test. B) Same as (A), but segregated by chromosomal arm.

**Supplementary Figure 13.**
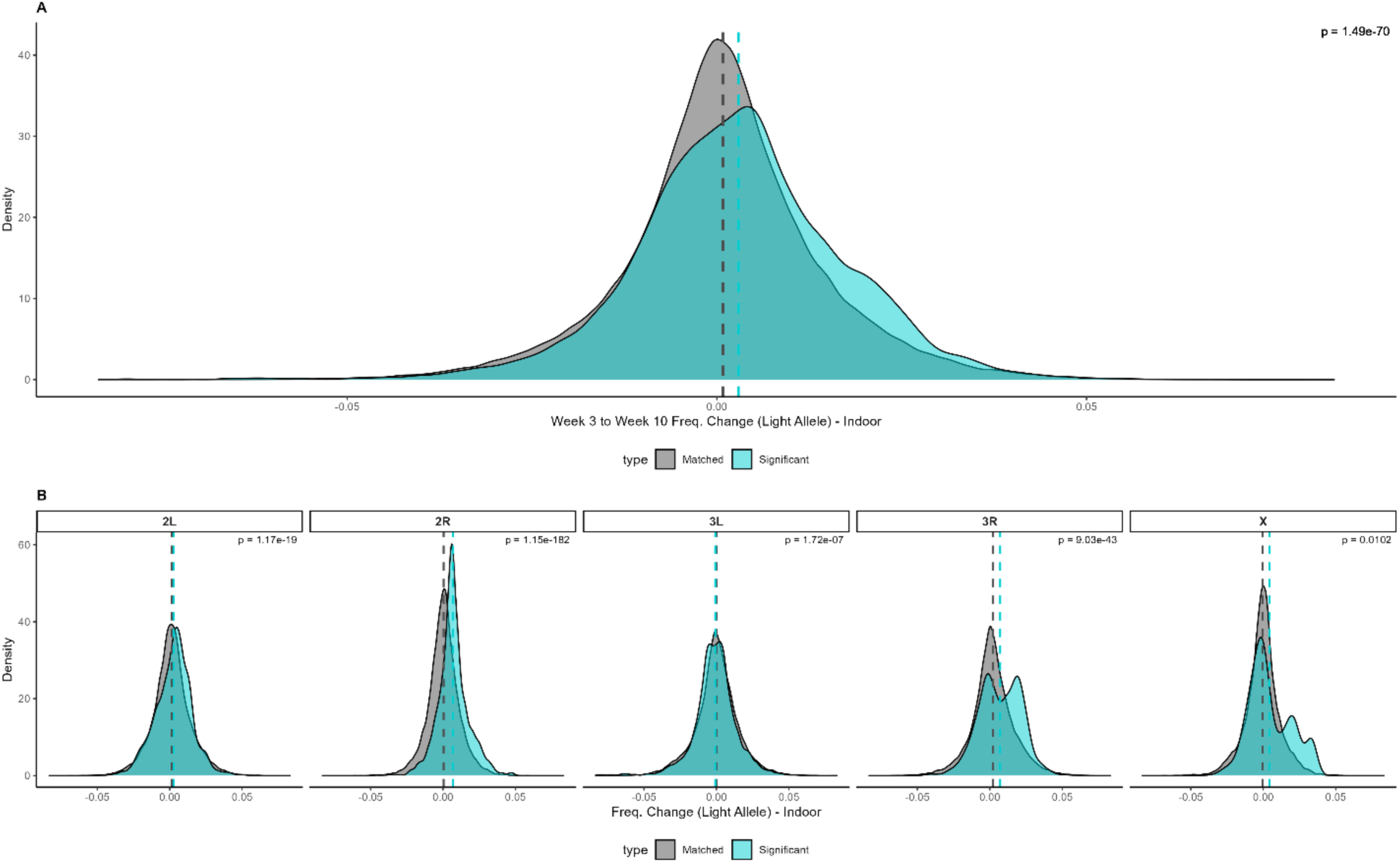
Seasonal shifts of trait-associated SNPs in indoor cages. A) Distribution of light-associated allele frequency shifts at all trait-associated SNPs (identified at Week 10 in outdoor cages) between Weeks 3 and 10 of evolution in the indoor lab environment. Distributions of frequency shifts at trait-associated SNPs are turquoise, while distributions derived from matched controls are grey. Reported p-values are derived from a comparison of trait-associated and matched control SNPs using a binomial test. B) Same as (A), but segregated by chromosomal arm.

**Supplementary Figure 14.**
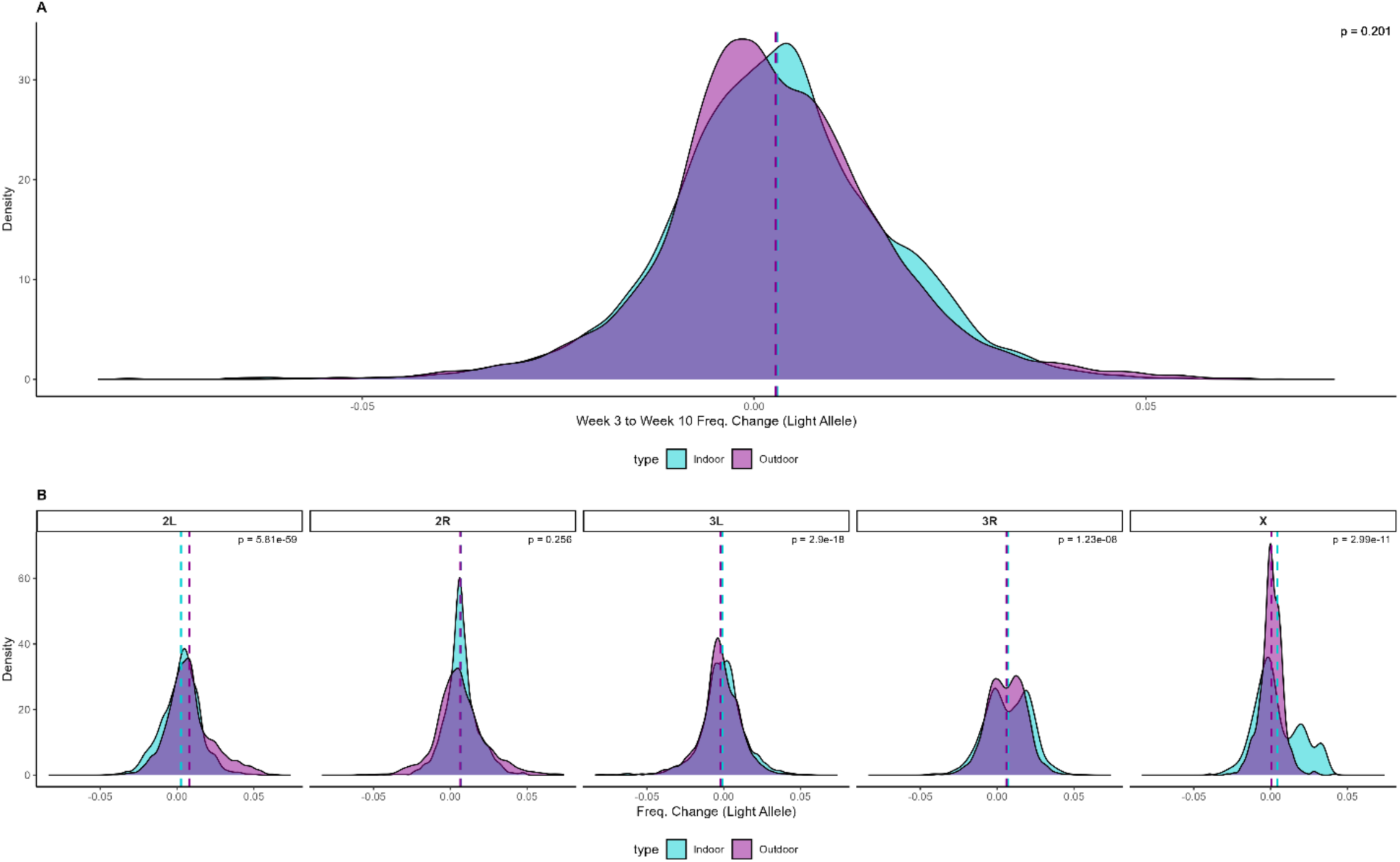
Seasonal shifts of trait-associated SNPs in outdoor versus indoor cages. A) Distribution of light-associated allele frequency shifts at all trait-associated SNPs (identified at Week 10 in outdoor cages) between Weeks 3 and 10 of evolution in the outdoor field vs. indoor lab environment across the whole genome. Distributions of frequency shifts at trait-associated SNPs in the outdoor cages are magenta, while distributions in indoor cages are turquoise. The indoor and outdoor distributions were compared using a two-sided t-test. B) Same as (A), but segregated by chromosomal arm.

**Supplementary Figure 15.**
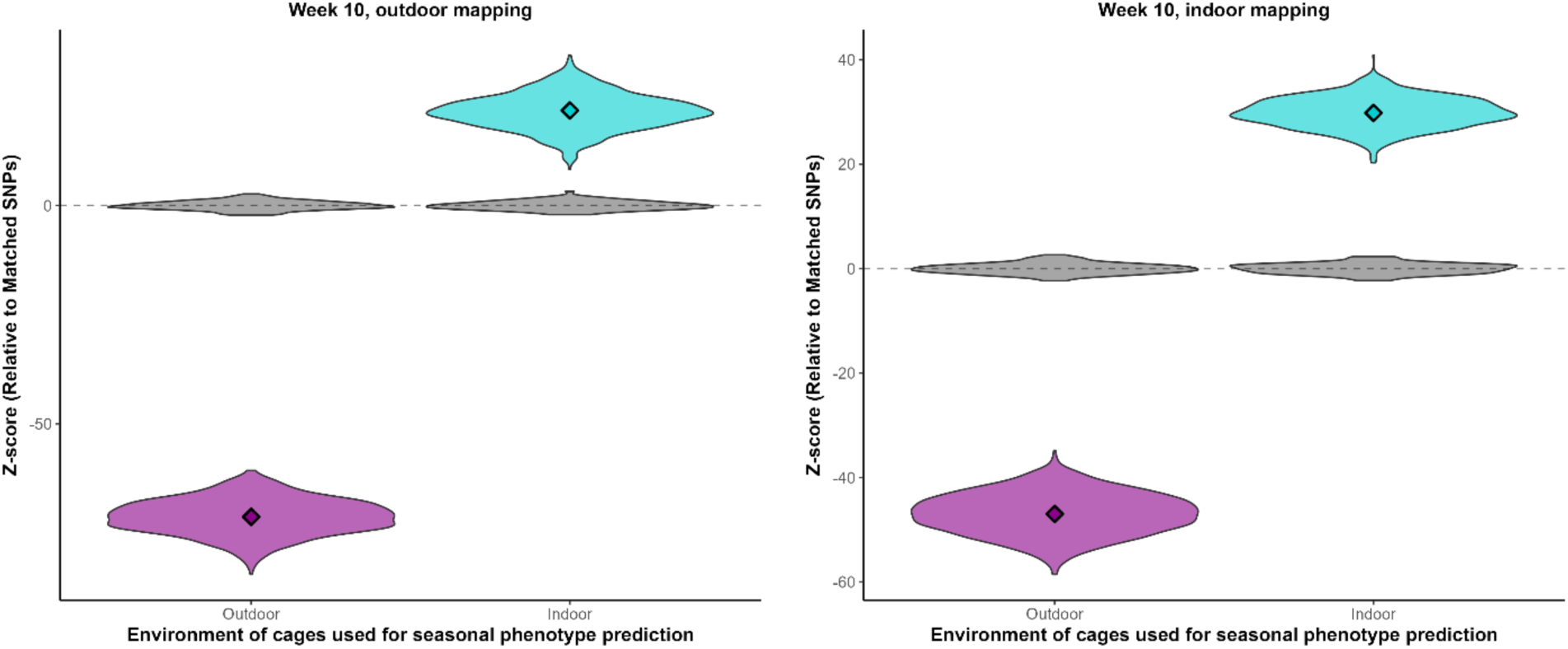
Phenotypic prediction with only *tan*, *ebony*, and *bab*, using mapping from each timepoint/treatment subset. Relative, predicted phenotypic change between Weeks 1 and 8 in both indoor and outdoor environments, based on only the three clusters that contain *tan, ebony*, or *bab*. Each quadrant uses a different mapping (for example, from Week 10 in the outdoor cages) and within each quadrant, we use the mapping to predict the phenotypic change in the outdoor environment (magenta) and the indoor environment (turquoise). The predicted change is denoted with diamonds, while the violins indicate the distribution of 1000 bootstrapped values (colored for trait-associated SNPs and grey to matched control SNPs).

**Supplementary Figure 16.**
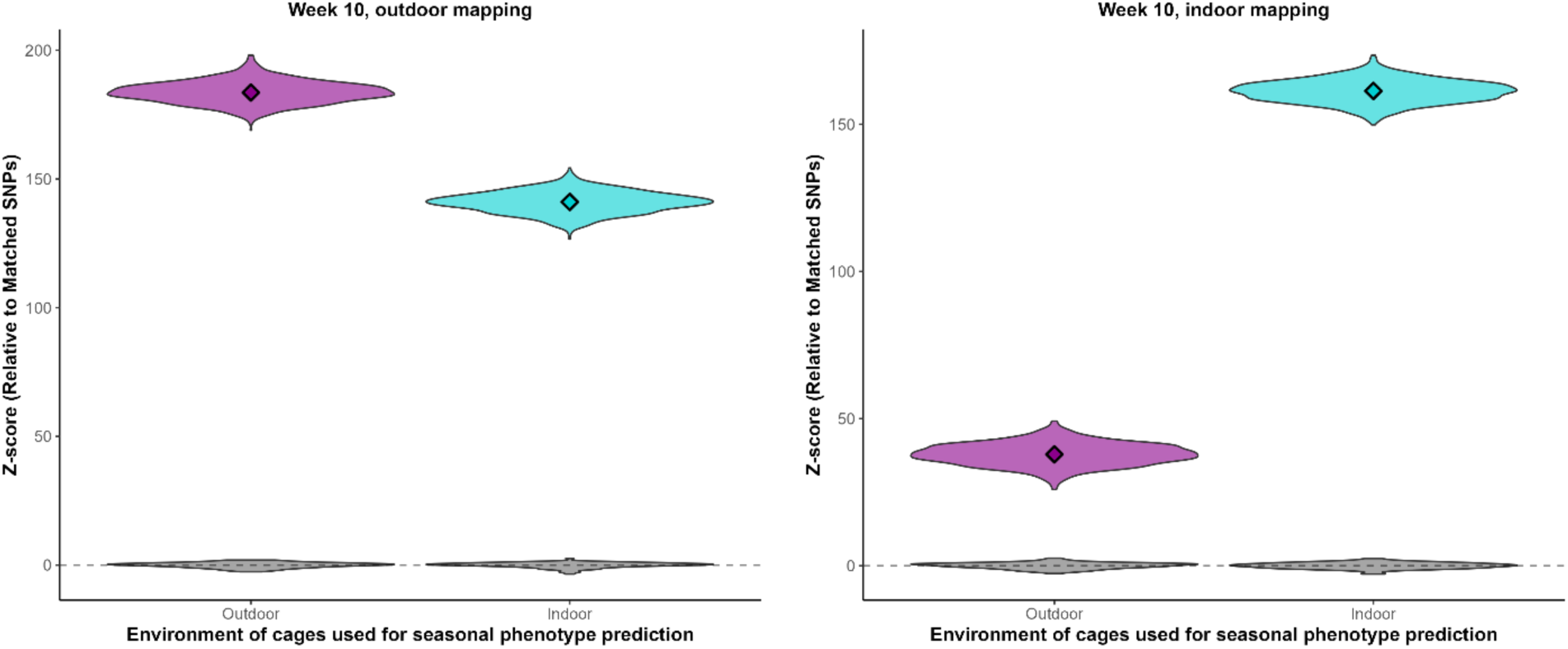
Phenotypic prediction excluding *tan*, *ebony*, and *bab*. Relative, predicted phenotypic change between Weeks 1 and 8 in both outdoor (magenta) and indoor (turquoise) environments, generated using our trait mapping results derived at weeks 8 in the outdoor (left) and indoor environment, and using variation from every mapped locus excluding those containing *tan, ebony*, or *bab*. Predicted phenotypic change are quantified as Z-scores (computed relative to matched control shifts) whereby positive Z-scores indicate net lighter pigmentation evolution while negative values would indicate net darker evolution. Z-scores were computed by comparing inferred relative phenotypic change using mapped SNPs to a that computed using a set of matched control SNPs (grey distributions).

**Supplementary Figure 17.**
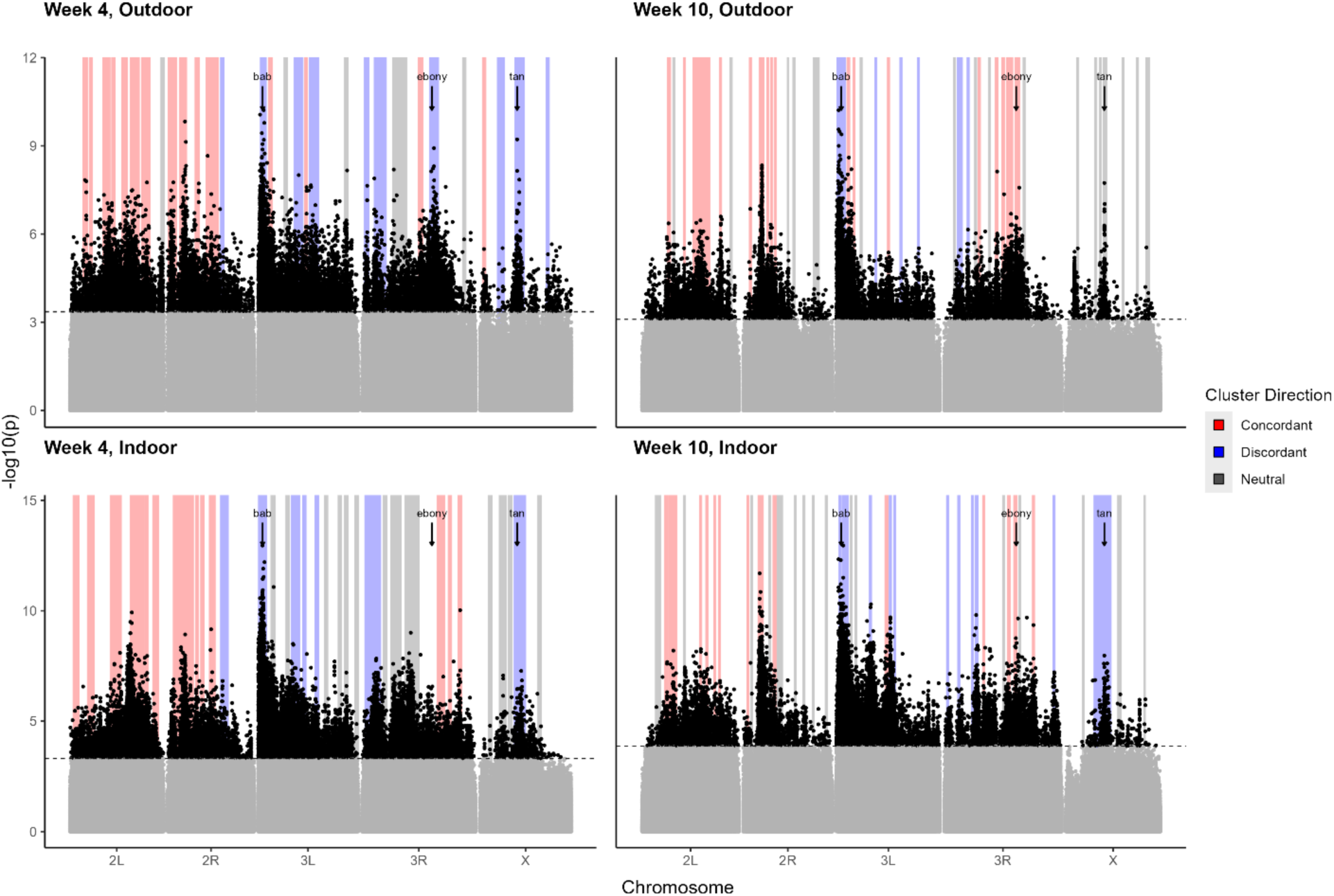
Manhattan plots produced across all mapping populations. Manhattan plot depicting nominal p-values derived from an ancestry-informed mapping of SNP association with pigmentation variation, separately for each mapping population (week 4 indoor, week 4 outdoor, week 10 indoor, week 10 outdoor). The dashed horizontal line corresponds to an empirical FDR threshold of 0.0005, derived via permutations. Shaded regions denote the stop and end points of each of the independent loci, while bar color denotes behavior of SNPs within each locus through time as the populations evolved lighter pigmentation within the outdoor mesocosm: red shading indicates loci with light-associated alleles favored throughout evolution of lighter pigmentation, blue shading indicates loci in which the light-associated alleles are predominately selected against and grey shading indicates loci with no signature of selection on light-associated alleles. The location of the canonical pigmentation genes, *bric-a-brac (bab), ebony,* and *tan*, are indicated on the Manhattan plots. Potential candidate genes underlying the remaining large effect loci observed throughout the genome are discussed in the Supplement.

**Supplementary Figure 18.**
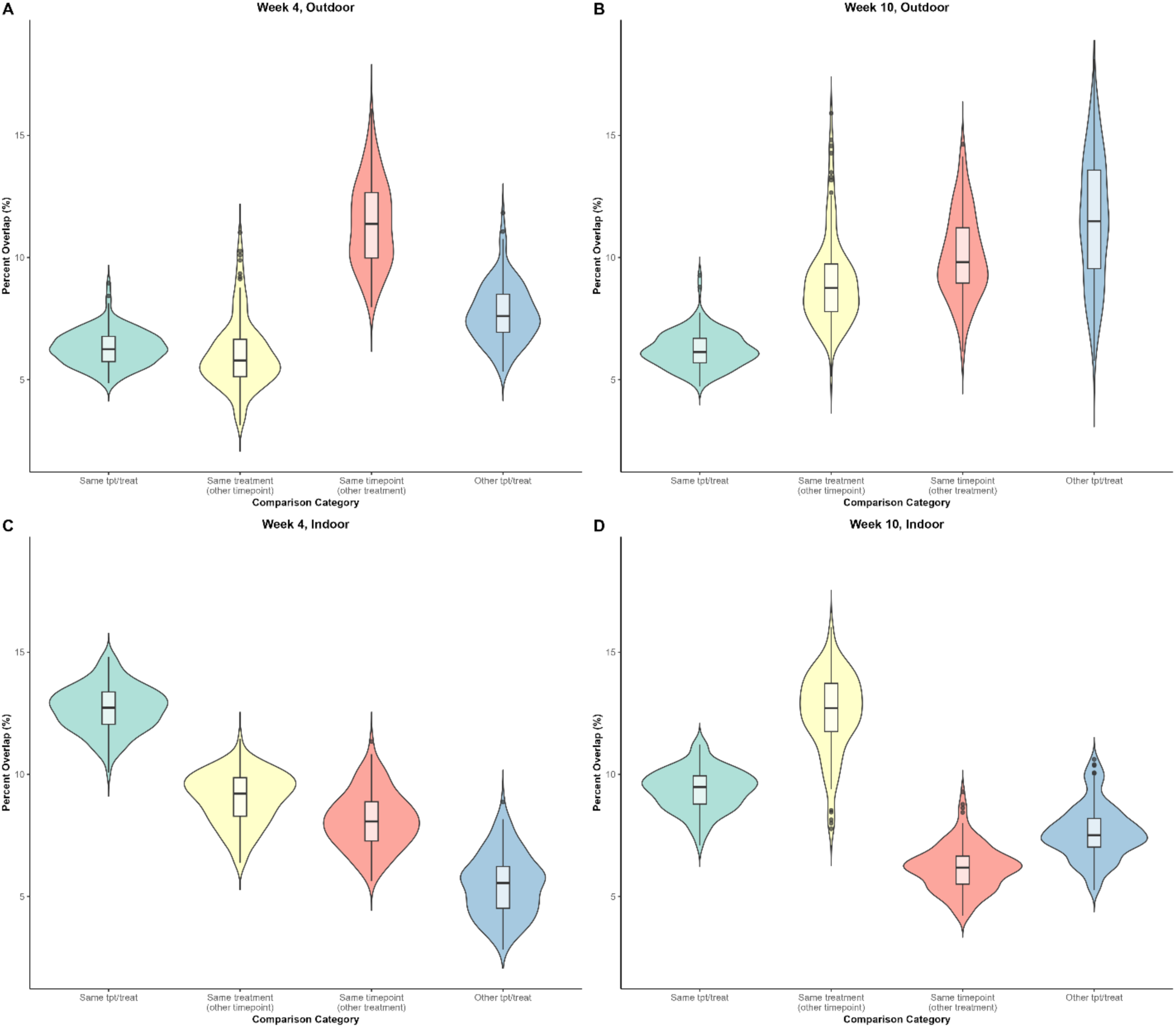
Shared significant SNPs across mapping populations. Overlap percentage from each pair of GLMs that each use five of the cages from a specific timepoint and treatment combination. Significant SNP overlap is either considered between GLMs using cages within the same timepoint/treatment (green), the same treatment but the other timepoint (yellow), the same timepoint but the other treatment (red), or cages from both the other timepoint and treatment (blue). The box plots represent the real percentage from each paired comparison, while the colored violins represent our bootstrapping results.

**Supplementary Figure 19.**
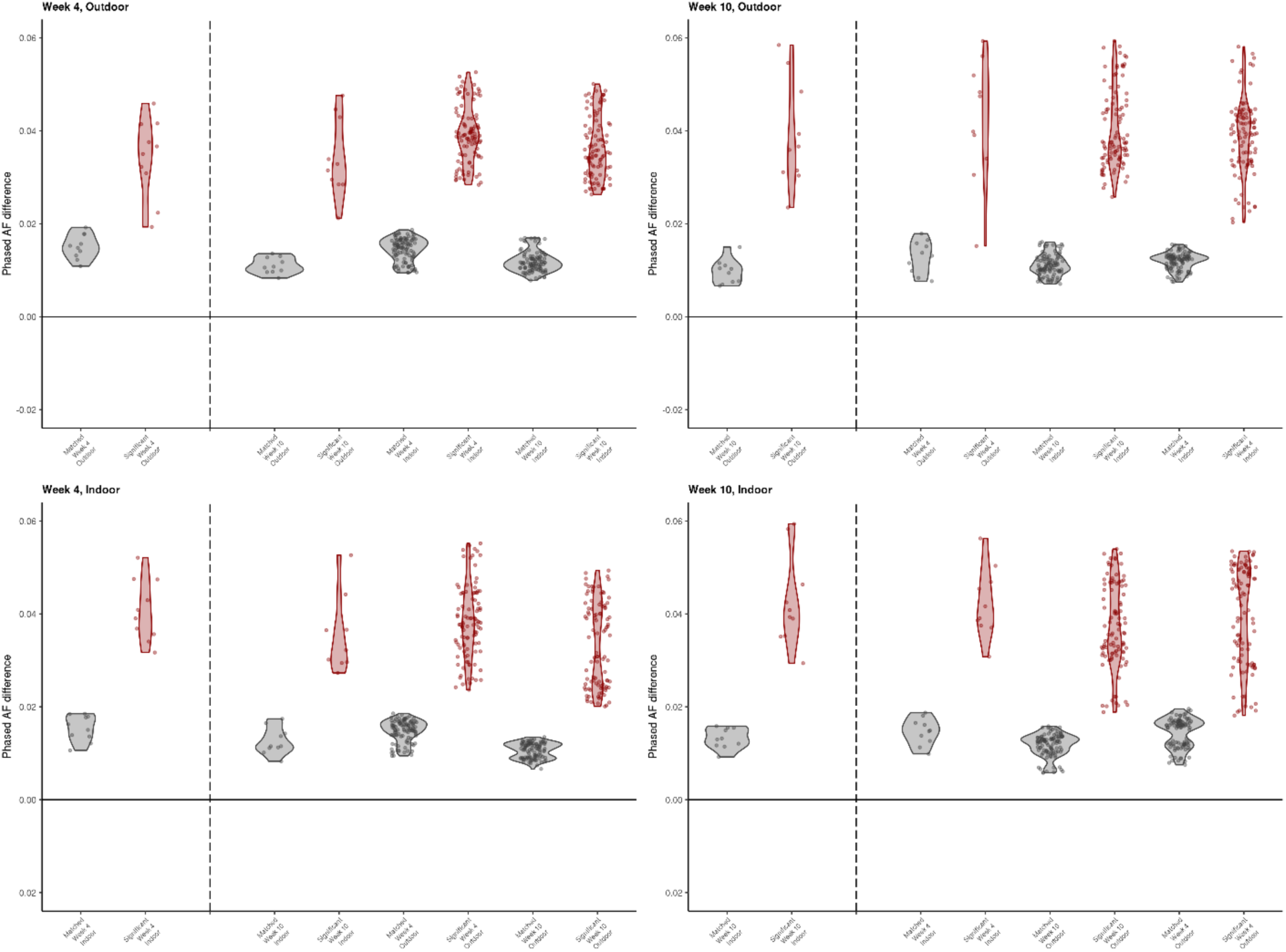
Leave-one-out across the full genome, for all timepoint/treatment combinations. Results from leave-one-out validation of mapping within and across treatments and time points for all timepoint/treatment combinations. Depicted are the median allele frequency difference for each test cage between light and dark samples of trait-associated SNPs (red), as well as a set of matched control SNPs (grey).

**Supplementary Figure 20.**
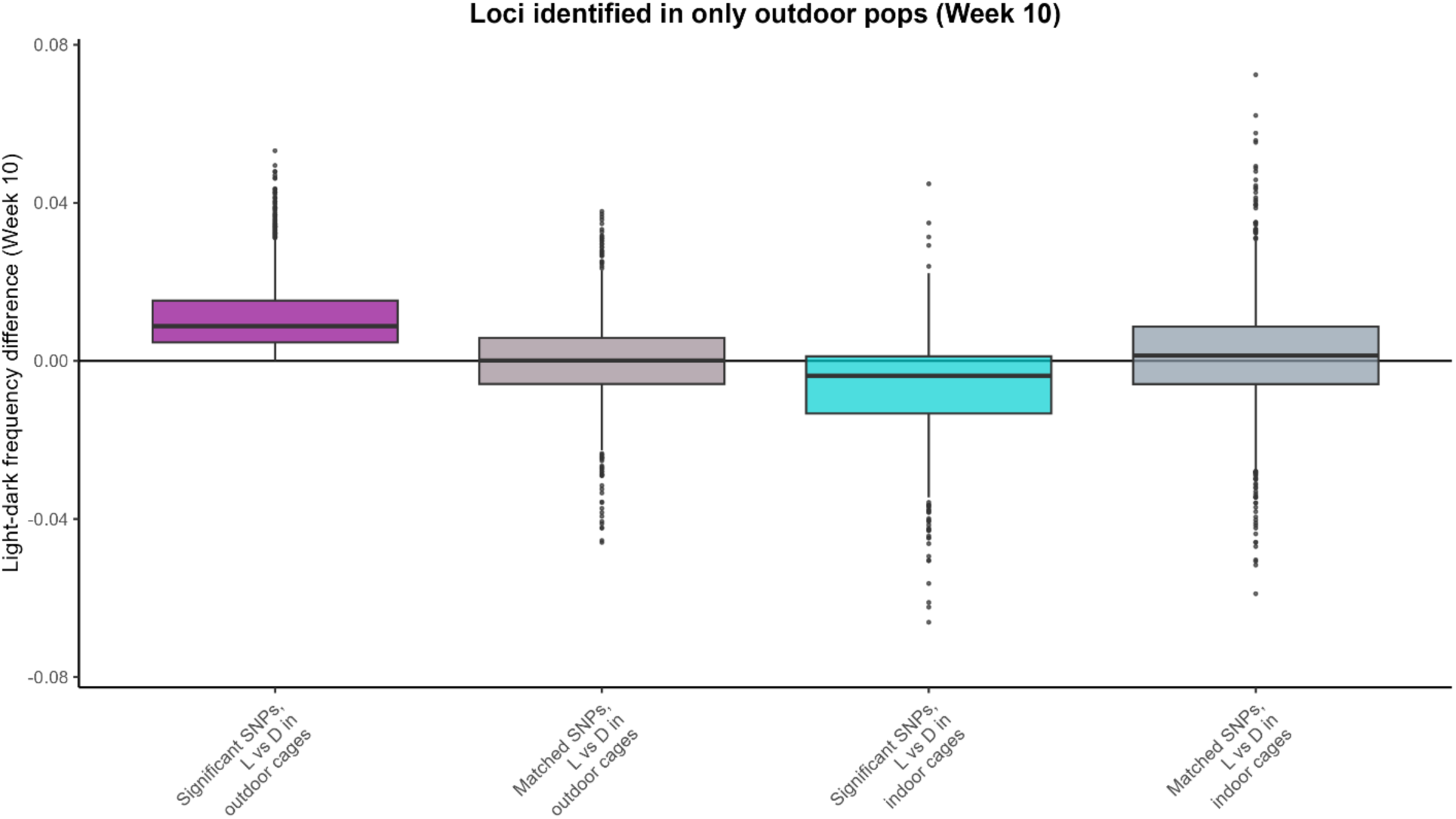
Evidence of field-specific pigmentation-associated loci. Distribution of light-dark allele frequencies (boxes denote the 50% interquartile range, dark horizontal lines correspond to the median, and points indicate outliers) for pigmentation associated SNPs located within mapped loci occurring the outdoor, week 10 mapping population that share no overlapping breakpoints with any loci in the indoor, week 10 mapping population. The magenta distribution corresponds to allele frequency differences within the outdoor cages, while the turquoise to the indoor environment. Grey distributions then represent the light-dark allele frequency differences at matched control SNPs for each environment. The light-dark frequency differences are significantly greater than the matched control shifts for the outdoor environment (Mann-Whitney U test p-value = 0.003), and significantly less than matched control SNPs for the indoor environment (Mann-Whitney U test p-value < 0.001). Furthermore, the outdoor and indoor frequency differences are significantly different than each other (Wilcoxon Signed Rank Test p-value < 0.001), indicating that light-associated alleles in outdoor environment are putatively enriched in the dark color fraction of the indoor environment for this set of loci.

**Supplementary Figure 21.**
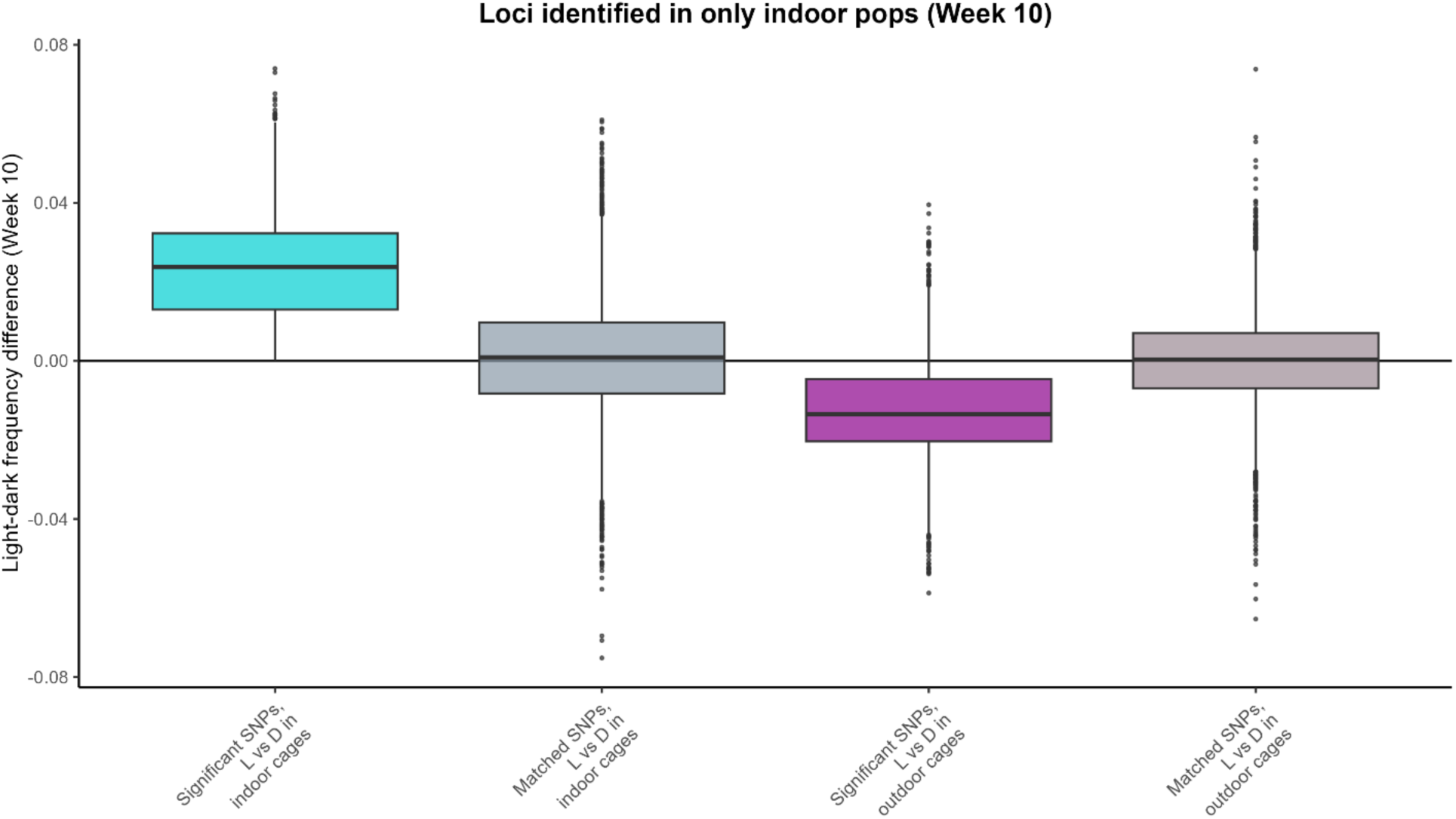
Evidence of lab-specific pigmentation-associated loci. Distribution of light-dark allele frequencies (boxes denote the 50% interquartile range, dark horizontal lines correspond to the median, and points indicate outliers) for pigmentation associated SNPs located within mapped loci occurring the indoor, week 10 mapping population that share no overlapping breakpoints with any loci in the outdoor, week 10 mapping population. The magenta distribution corresponds to allele frequency differences within the outdoor cages, while the turquoise to the indoor environment. Grey distributions then represent the light-dark allele frequency differences at matched control SNPs for each environment. The light-dark frequency differences are significantly greater than the matched control shifts for the indoor environment (Mann-Whitney U test p-value < 0.001), and significantly less than matched control SNPs for the outdoor environment (Mann-Whitney U test p-value < 0.001). Furthermore, the indoor and outdoor frequency differences are significantly different than each other (Wilcoxon Signed Rank Test p-value < 0.001), indicating that light-associated alleles in indoor environment are putatively enriched in the dark color fraction of the outdoor environment for this set of loci.

**Supplementary Figure 22.**
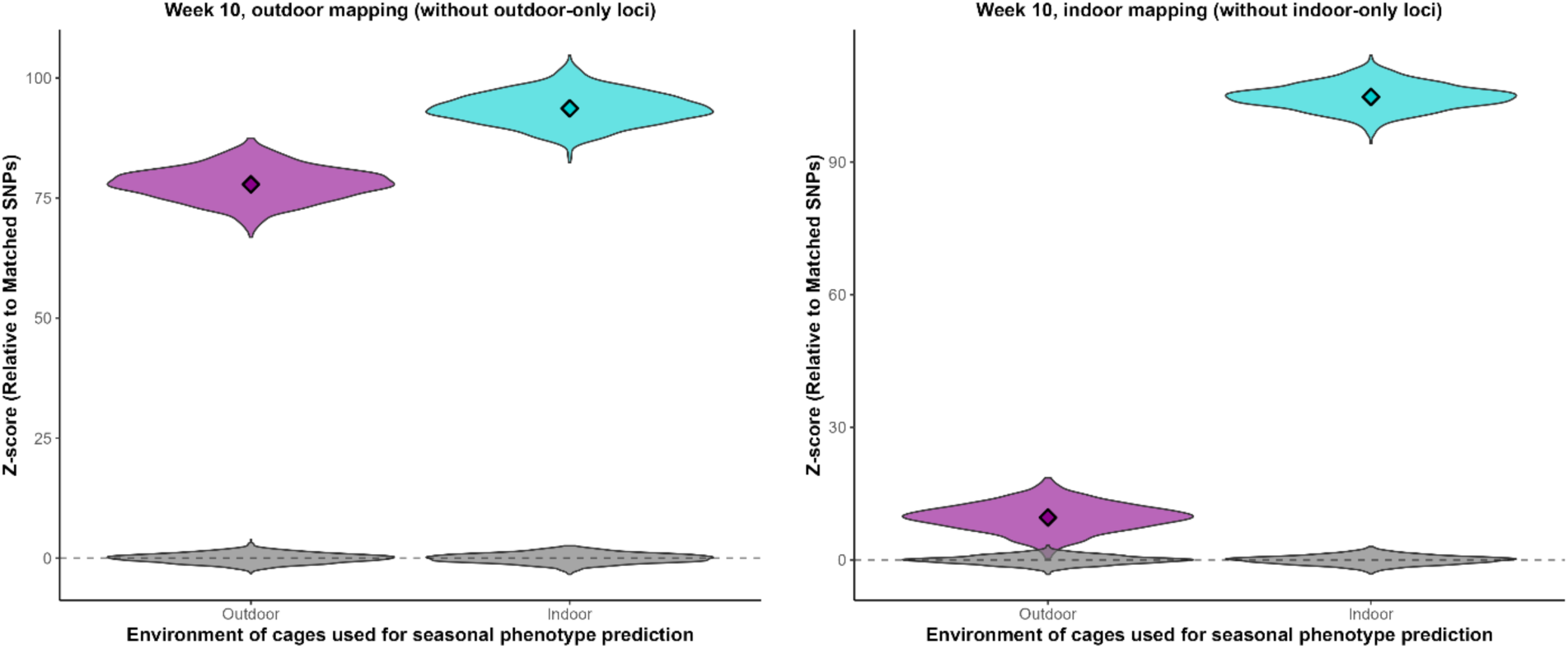
Phenotypic prediction excluding treatment-specific loci. Relative, predicted phenotypic change between Weeks 1 and 8 in both outdoor (magenta) and indoor (turquoise) environments, generated using our trait mapping results derived at weeks 8 in the outdoor (left) and indoor environment, and using variation from every mapped locus excluding those that are identified only in the outdoor cages at week 10 or identified only in the indoor cages at week 10. Predicted phenotypic change are quantified as Z-scores (computed relative to matched control shifts) whereby positive Z-scores indicate net lighter pigmentation evolution while negative values would indicate net darker evolution. Z-scores were computed by comparing inferred relative phenotypic change using mapped SNPs to a that computed using a set of matched control SNPs (grey distributions).

**Supplementary Figure 23.**
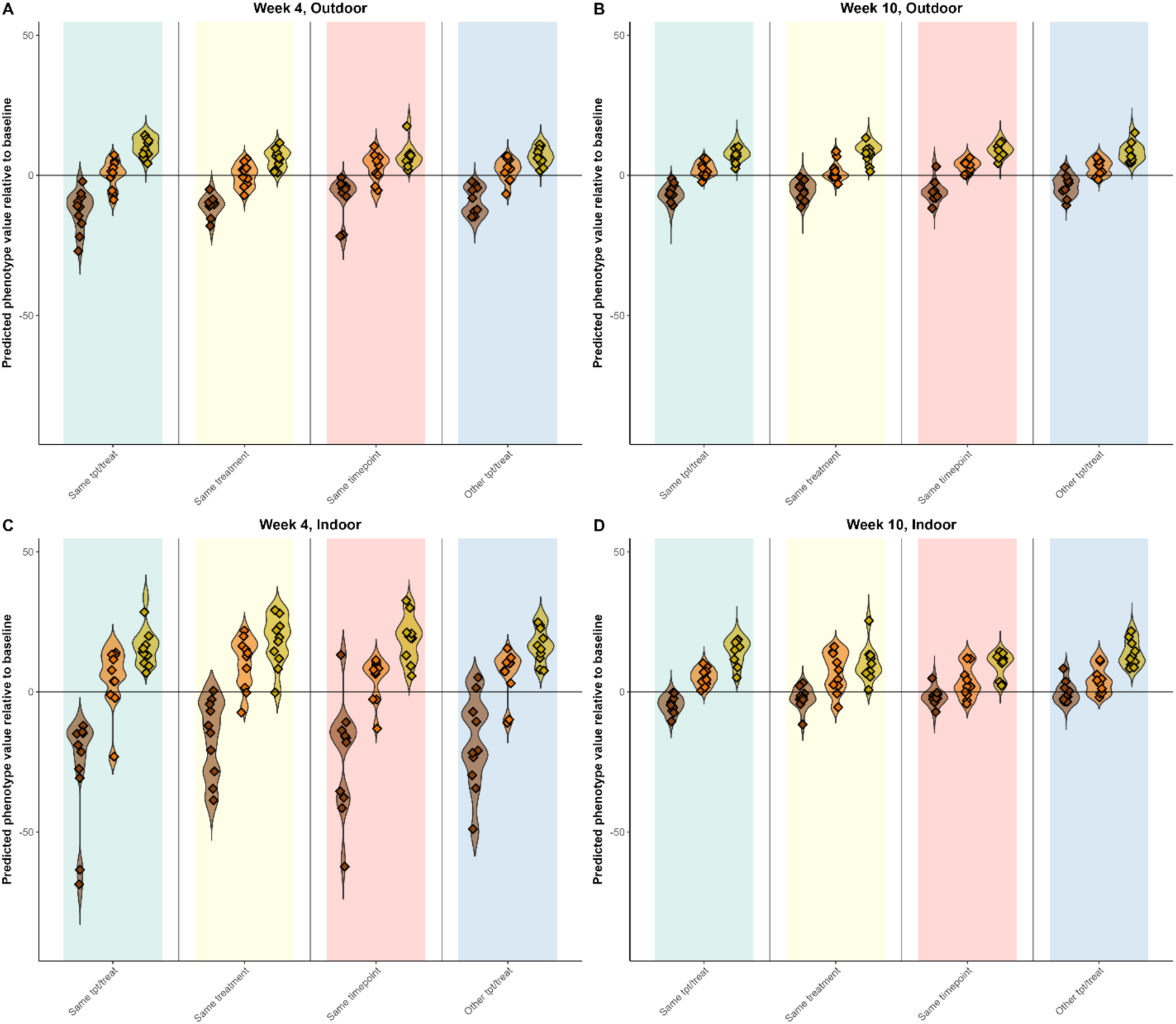
Phenotypic predictions of relative melanization across timepoint and treatment. Using the mapping from each timepoint/treatment subset and considering all significant SNPs, we generate a phenotypic prediction score for each color fraction sample. A negative value indicates an expected phenotypic shift towards dark pigmentation and a positive value a shift towards lighter pigmentation. We consider our prediction “correct” when the majority of dark samples (dark brown) have a negative predicted value, the majority of light samples (gold) have a positive one, and a linear regression (where we encode dark = 0, midpoint = 1, and light = 2) is significant and has a positive slope. We predict the phenotype of all samples from either within the same timepoint/treatment (green), the same treatment but the other timepoint (yellow), the same timepoint but the other treatment (red), or from both the other timepoint and treatment (blue).

**Supplementary Figure 24.**
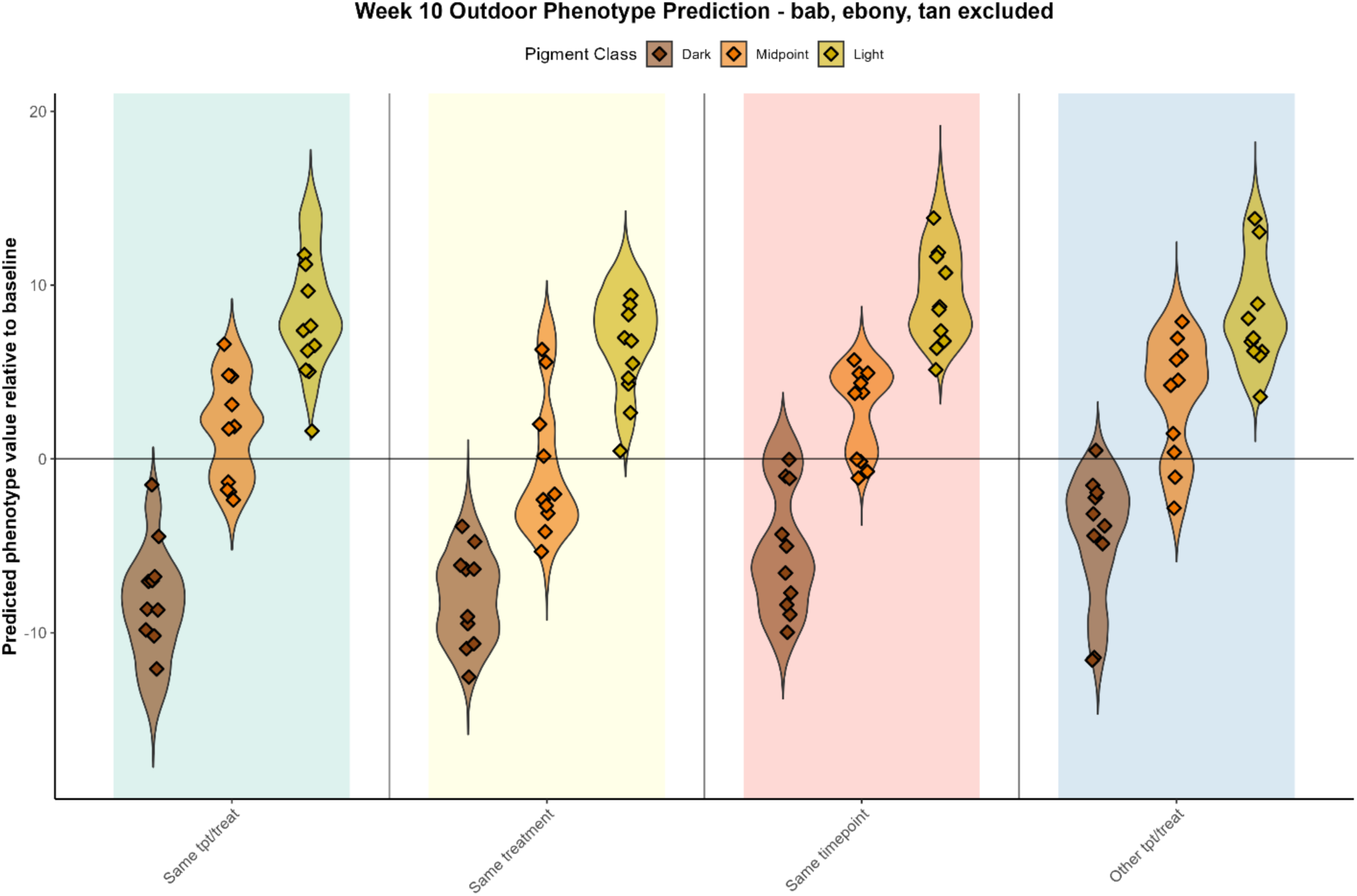
Phenotypic predictions of color samples across timepoint and treatment, excluding variation in canonical pigmentation loci. Using the mapping from the outdoor, week 10 mapping population, and considering all significant SNPs outside the loci containing the canonical pigmentation genes (*tan, ebony,* and *bric-a-brac*), we generate a phenotypic prediction score for each color fraction sample. A negative value indicates an expected phenotypic shift towards dark pigmentation and a positive value a shift towards lighter pigmentation. We consider our prediction “correct” when the majority of dark samples (dark brown) have a negative predicted value, the majority of light samples (gold) have a positive one, and a linear regression (where we encode dark = 0, midpoint = 1, and light = 2) is significant and has a positive slope. We predict the phenotype of all samples from either within the same timepoint/treatment (green), the same treatment but the other timepoint (yellow), the same timepoint but the other treatment (red), or from both the other timepoint and treatment (blue).

**Supplementary Figure 25.**
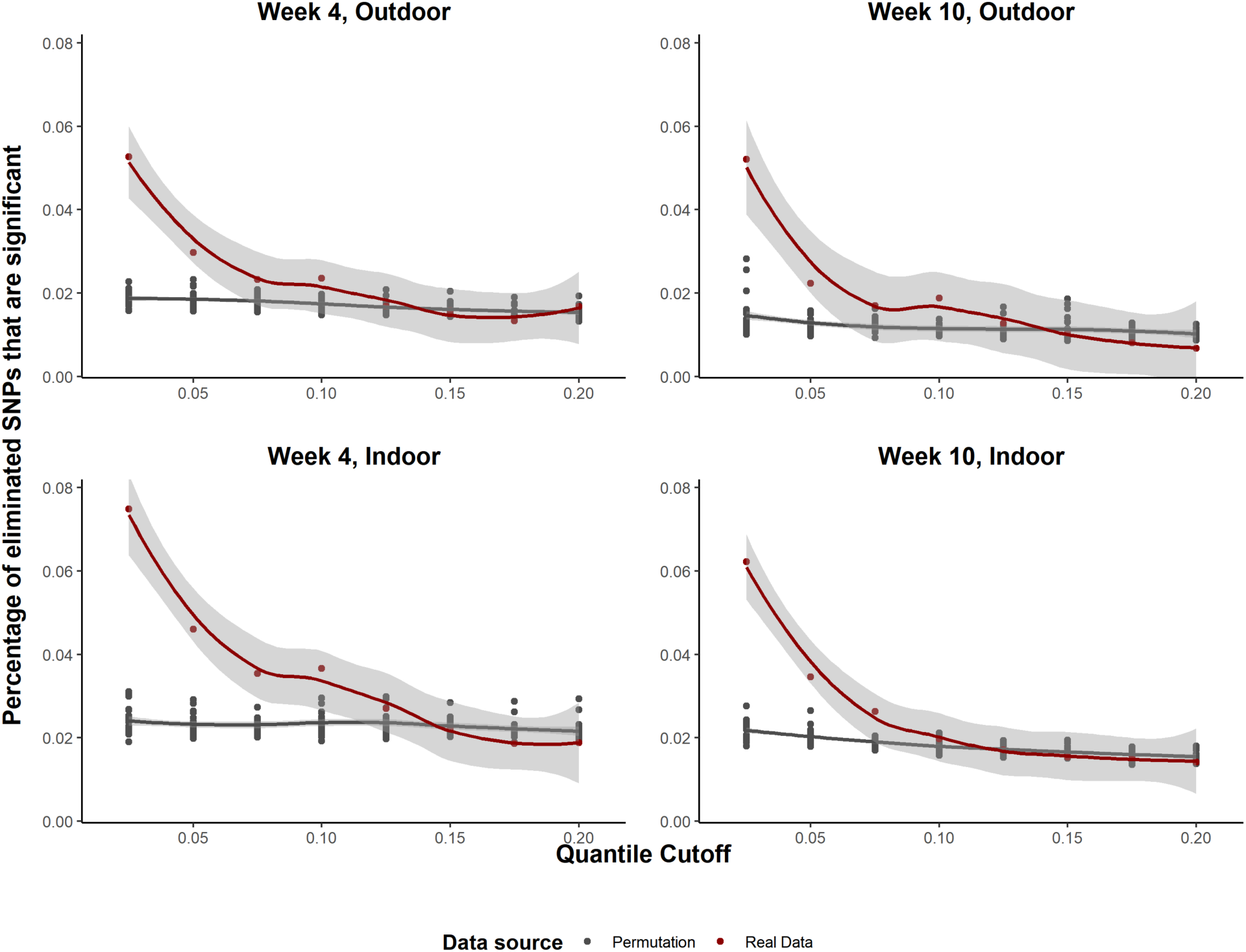
Percentage of top SNPs eliminated in first round of unlinked locus identification with different quantile cutoffs. In our clustering method, the SNPs with the highest light-dark difference as predicted by haplotype frequencies in the highest signal region are eliminated sequentially. Here, we look at the percentage of the eliminated SNPs that are significant SNPs in the first round of clustering with cutoffs at different quantiles using real data versus permutations.

**Supplementary Figure 26.**
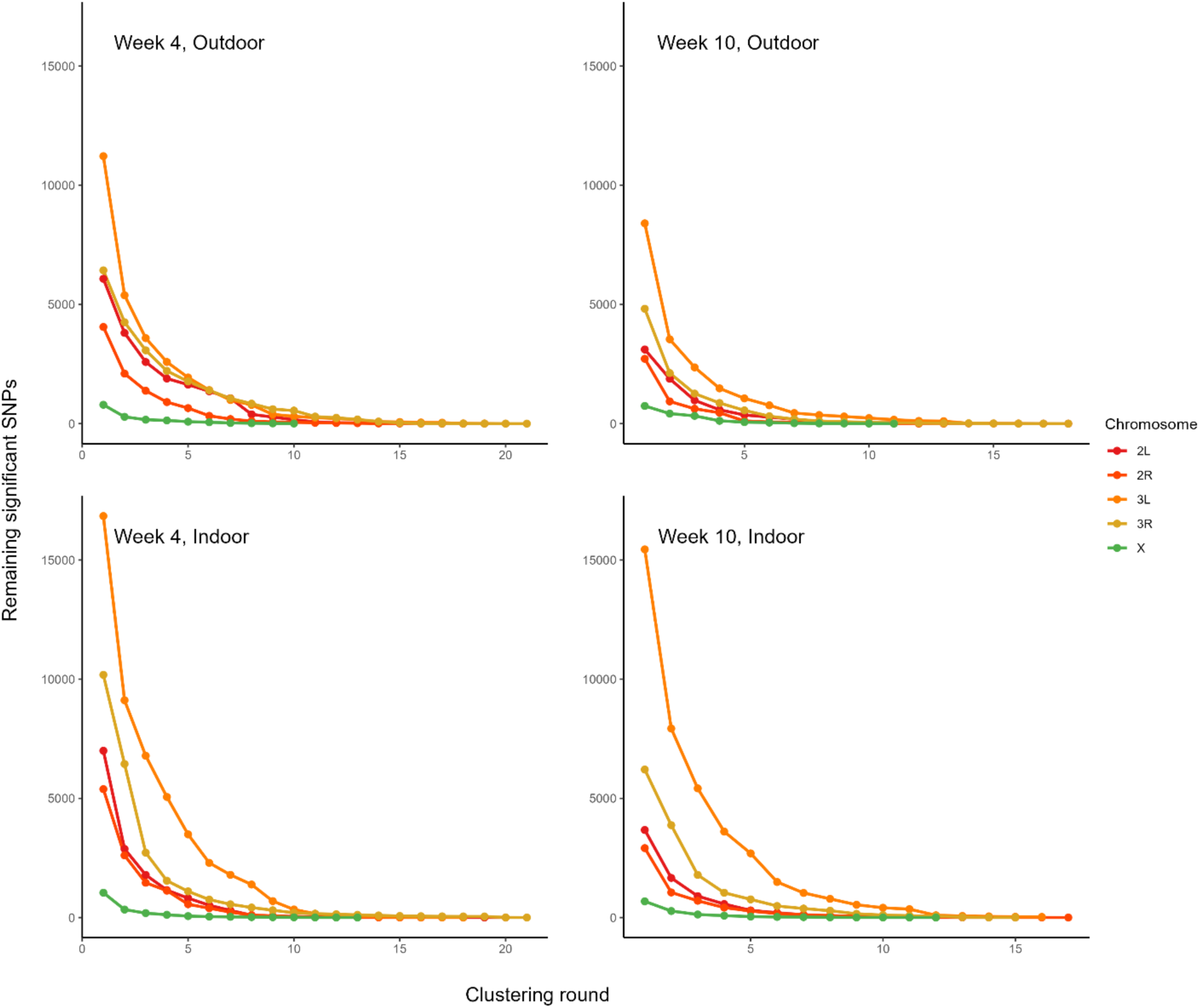
Signal elimination throughout clustering. Plot of the number of remaining significant SNPs before each round of clustering for each chromosome, showing the elimination of linked significant SNPs throughout the clustering process.

### Supplementary Tables

**Table S1.**
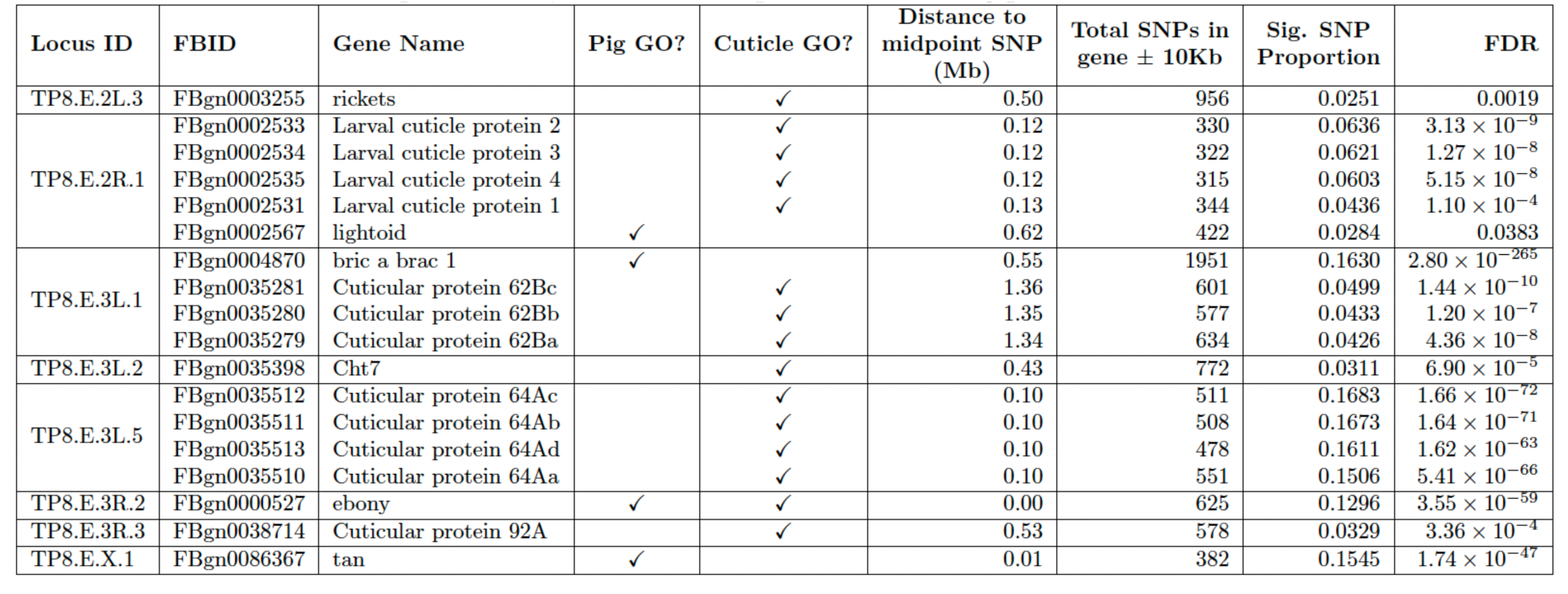
List of pigmentation (Pig) and cuticle-associated genes located within mapped loci identified in the week 10, outdoor mapping population that are enriched in pigmentation-associated, mapped SNPs (binomial test, FDR < 0.05). Also reported is the locus ID (corresponding to the locus information provided in Supplementary Data File 1) within which each gene resides, the distance of each gene to the locus midpoint SNP, the number of SNPs within or within 10 Kb of the gene, the proportion of SNPs within the gene that are pigmentation associated (Sig. SNP proportion), and the Benjamini-Yekutieli false discovery rate corrected p-value for observed enrichment of pigmentation-associated SNPs.

**Table S2.** List of pigmentation (Pig) and cuticle-associated genes located within mapped loci identified in the week 10, indoor mapping population that are enriched in pigmentation-associated, mapped SNPs (binomial test, FDR < 0.05). Also reported is the locus ID (corresponding to the locus information provided in Supplementary Data File 1) within which each gene resides, the distance of each gene to the locus midpoint SNP, the number of SNPs within or within 10 Kb of the gene, the proportion of SNPs within the gene that are pigmentation associated (Sig. SNP proportion), and the Benjamini-Yekutieli false discovery rate corrected p-value for observed enrichment of pigmentation-associated SNPs.

| Locus ID | FBID | Gene Name | Pig GO? | Cuticle GO? | Distance to midpoint SNP (Mb) | Total SNPs in gene ± 10Kb | Sig. SNP Proportion | FDR |
| --- | --- | --- | --- | --- | --- | --- | --- | --- |
| TP8.I.2L.3 | FBgn0053302 | Cuticular protein 31A | | ✓ | 0.03 | 642 | 0.0685 | $1.05 \times 10^{-14}$ |
| TP8.I.2R.1 | FBgn0002534 | Larval cuticle protein 3 | | ✓ | 0.43 | 322 | 0.1770 | $1.58 \times 10^{-40}$ |
| | FBgn0002535 | Larval cuticle protein 4 | | ✓ | 0.44 | 315 | 0.1619 | $3.09 \times 10^{-34}$ |
| | FBgn0002533 | Larval cuticle protein 2 | | ✓ | 0.43 | 330 | 0.1485 | $4.15 \times 10^{-31}$ |
| | FBgn0002531 | Larval cuticle protein 1 | | ✓ | 0.43 | 344 | 0.1279 | $3.87 \times 10^{-25}$ |
| TP8.I.3L.1 | FBgn0000543 | ecdysoneless | | ✓ | 0.69 | 535 | 0.0897 | $1.08 \times 10^{-20}$ |
| | FBgn0035279 | Cuticular protein 62Ba | | ✓ | 0.25 | 634 | 0.0757 | $1.08 \times 10^{-17}$ |
| | FBgn0035280 | Cuticular protein 62Bb | | ✓ | 0.26 | 577 | 0.0676 | $7.67 \times 10^{-13}$ |
| | FBgn0022702 | Chitinase 2 | | ✓ | 0.18 | 350 | 0.0657 | $2.17 \times 10^{-7}$ |
| | FBgn0035281 | Cuticular protein 62Bc | | ✓ | 0.27 | 601 | 0.0416 | $1.74 \times 10^{-4}$ |
| TP8.I.3L.2 | FBgn0035875 | Cuticular protein 66Cb | | ✓ | 0.30 | 429 | 0.0559 | $1.82 \times 10^{-6}$ |
| TP8.I.3L.5 | FBgn0004870 | bric a brac 1 | ✓ | | 0.33 | 1951 | 0.1978 | $1.44 \times 10^{-295}$ |
| TP8.I.3L.6 | FBgn0261797 | Dynein heavy chain 64C | | ✓ | 0.23 | 782 | 0.0665 | $7.88 \times 10^{-17}$ |
| TP8.I.3R.2 | FBgn0027508 | tankyrase | | ✓ | 0.34 | 537 | 0.0503 | $2.38 \times 10^{-6}$ |
| TP8.I.3R.3 | FBgn0000527 | ebony | ✓ | ✓ | 0.07 | 625 | 0.0432 | $3.97 \times 10^{-5}$ |
|  | FBgn0038819 | Cuticular protein 92F |  | ✓ | 0.49 | 464 | 0.0409 | 0.0024 |
| TP8.I.3R.6 | FBgn0038901 | Bursicon | ✓ | ✓ | 0.36 | 325 | 0.0554 | $8.00 \times 10^{-5}$ |
| TP8.I.X.1 | FBgn0086367 | tan | ✓ | | 0.01 | 382 | 0.1440 | $3.10 \times 10^{-34}$ |

### Supplementary Discussion

#### Novel pigmentation-associated loci identified by high-powered, tail-based mapping

Here we describe in further detail genes occurring within our mapped loci that likely play a functional role in pigmentation variation. As nearly 80% of total genes in the genome were located within the start and endpoints of the loci identified across all mapping populations, we concentrate this discussion on those genes with a gene ontology (GO) term associated with pigmentation (excluding eye-specific pigmentation GO terms) and/or with cuticle development and molting, as well as confine our assessment to mapping conducted at week 10. Additionally, for each putative candidate gene, we ensured that the proportion of mapping significant SNPs within or nearby (within 10 Kb) gene breakpoints were significantly greater than expected based on the genome-wide distribution of significant SNPs (binomial test).

As expected, *bric-a-brac* (*bab*), *ebony*, and *tan* each fell within mapped loci and contained a significant enrichment of pigmentation-associated SNPs. *Ebony* and *tan* are opposing enzymes that catalyze dopamine to a precursor of sclerotin (N-β-alanyl-dopamine, NBAD) and NBAD back to dopamine, respectively ^1^. Thus, together they control the amount of NBAD available to create light pigmented sclerotin instead of darkly pigmented melanin. In contrast, *bab* is a set of two duplicated transcription factors (*bab1* and *bab2*) that repress pigmentation ^2^. For example, *bab* is expressed in tergites A2 through A6 in female *D. melanogaster,* but only A2 through A4 in males, giving rise to the characteristic, male-only, completely pigmented terminal tergites for which *D. melanogaster* is named ^3^. Binding sites for *Abdominal-B* (*Abd-B*) and *doublesex* (dsx) are located in the long intron of bab1, and the binding of *doublesex* to *bab1* mediates this sex-specific pigmentation ^4^. Additionally, previous studies have shown that the regulation of both *bab* and *tan* is influenced by temperature, providing a likely explanation for the plasticity of pigmentation across developmental temperatures observed in *D. melanogaster* ^5,6^. For *bab*, the *hox* gene and transcription factor Abd-B has been shown to mediate this temperature-dependent expression ^7^. We also identify Abd-B in our study which is located within the top locus on chromosomal arm 3R.

On 2R, our largest peak (which shows high consistency across timepoints and treatments), overlies larval cuticle proteins 1, 2, and 3 (*Lcp1*, *Lcp2*, *Lcp3*) – three out of four larval cuticle protein genes that form a set of two pairs of genes (*Lcp1* and *Lcp2*, and *Lcp3* and *Lcp4*) that each act as a separate transcriptional unit and are transcribed at different periods during the third instar stage ^8^. *Lcp3* and *Lcp4* accumulate first, followed by *Lcp1* and *Lcp2*, and this is likely part of a set of changes that shift the third instar soft cuticle to a harder, less flexible one for the wandering stage ^8^. Even though the larval cuticle is not preserved into the adult stage, certain pigmentation genes (namely yellow) that are important for adult pigmentation are also expressed and influence pigmentation in the larval stages ^9^.

We find further evidence of pigmentation-associated SNPs in loci concentrated in and around *Bursicon* (*Burs*), *rickets* (*rk*), and *Partner of bursicon* (*Pburs*). These three genes are the core genes vital to the cuticular hardening after eclosion. *Burs* and *Pburs* dimerize and bind to rickets to initialize the final tanning or hardening of the cuticle ^10,11^. Prior to this, the eclosed fly is entirely white, with very little visible pigmentation. Tanning then occurs when deposited cuticular proteins are linked together and the pigmentation becomes stark and visible. Despite this direct connection to pigmentation of the cuticle, this mapping study represents the first to directly identify pigmentation-relevant variation in these genes.

In only the outdoor week 10 mapping population, we find more significant SNPs than expected around and within a gene on chromosomal arm 3L that encodes for a transmembrane chitinase, *Chitinase 7* (*Cht7*) ^12^. It appears that *Cht7* is primarily involved in chitin organization in the developing cuticle, as RNAi knockdown of *Cht7* has shown the old cuticle to be disassembled successfully, but the density and organization of the new cuticle is compromised ^13^. Another chitinase gene, *Chitinase 2* (*Cht2*), is nearby *Cht7* on 3L, but only exhibited a significant enrichment of significant SNPs in the indoor mapping population. *Cht2*, like *Cht7*, seems to play a role in organizing the developing cuticle and not in disassembling the old cuticle ^14^.

In the indoor week 10 mapping population, we identified loci with high signal in and around *tankyrase*, *Dynein heavy chain 64c* (*Dhc64c*), and *ecdysoneless*. Both *tankyrase* and *Dhc64c* are involved in the proper expression and localization of *Wg* ^15,16^. *Wg* and other components in, or affected by, the *Wnt* pathway are known to be associated with pigmentation differences within and between species ^17^. *Ecdysoneless*, while it does affect the production of ecdysone as its name implies, actually has a broader role in correctly localizing pre-mRNA processing factor 8 (*Prp8*), a highly conserved splicing factor ^18,19^.

